# Piezophilic activity dominates the Puerto Rico Trench deep subseafloor biosphere

**DOI:** 10.64898/2026.09.30.755482

**Authors:** Miguel Desmarais, Corinna Breusing, Anna H. Finch, Chloé M. J. Baumas, William D. Orsi, Alvaro M. Plominsky, Yuki Morono, Laura M. Wehrmann, Dennis Graham, Robert A. Pockalny, Arthur J. Spivack, Irita Aylward, Taylor Walton, Barbara Saucedo, Sofía Rivera Soto, Jelis Sostre-Cortes, Ying Zhang, Evan A. Solomon, David C. Smith, Anne E. Dekas, Jeffrey S. Bowman, Steven D’Hondt, Douglas H. Bartlett

## Abstract

Hydrostatic pressure defines the deep ocean, yet its role in structuring microbial activity across the subseafloor biosphere remains poorly understood. Here, we assessed pressure effects on anabolic activity in sediment communities across margin-proximal, bathyal, abyssal, and hadal settings of the Puerto Rico Trench (493–8,385 m water depth). Community-level responses to pressure reversed with burial, revealing an unrecognized shift toward piezophily in communities down to 20 meters below seafloor (mbsf). Atmospheric pressure yielded higher proportions of active cells in surficial sediments, whereas *in situ* pressure yielded higher proportions in the subseafloor. Pressure also structured active-community composition; in trench-axis sediments down to 8.35 mbsf, JS1/Atribacterota populations within widespread hadal lineages exhibited a physiological response to *in situ* pressure, indicative of subseafloor piezophily. These findings suggest that hydrostatic pressure acts as an ecological filter after burial, favoring piezophilic traits that may support microbial persistence in the deep subseafloor biosphere.

---

The subseafloor biosphere hosts microbial life extending kilometers below the seafloor^1,2^. Although cell abundance and metabolic rates generally decline with burial, microorganisms remain active under severe energy limitation over geological timescales^1,3,4^. Hydrostatic pressure increases with water depth by approximately 1 MPa per 100 m, reaching 110 MPa (>1,000 atmospheres) in the deepest trenches, constraining microbial physiology and activity throughout the deep ocean^5–9^. Yet pressure is frequently excluded from experimental studies of the subseafloor biosphere owing to the technical challenges of conducting experiments under *in situ* pressure^10^.

Pressure has been shown to structure microbial community composition and diverse biogeochemical processes in deep-ocean environments, including organic matter degradation and carbon, nitrogen, and sulfur cycling^5,10–15^. In surficial deep-sea sediments, community-level responses to pressure range from piezosensitive responses, characterized by lower microbial activity when exposed to pressures above atmospheric pressure (*i.e.*, >0.1 MPa), to piezophilic responses, characterized by higher microbial activity when exposed to elevated pressure (>10 MPa)^8,9^. Early experiments with surficial Puerto Rico Trench-axis sediments showed strong piezosensitivity, with bulk heterotrophic activity suppressed under *in situ* pressure^16^. In contrast, activity in abyssal Atlantic sediments was maintained or enhanced, indicating pressure tolerance or piezophily^17,18^. Below the sediment surface, however, evidence for piezophily is largely limited to a small number of microorganisms isolated from deep subseafloor sediments ^19–21^. Community-level responses to pressure after burial remain largely unknown, including the extent to which pressure acts as an ecological filter shaping microbial persistence over geological timescales.

Hadal trenches provide a natural setting in which to address this knowledge gap. Their steep bathymetry creates pronounced hydrostatic-pressure gradients across relatively short geographic distances. Topographic focusing and episodic mass wasting also generate heterogeneous depositional environments and sedimentary records^22–24^. These gradients can extend from margin-proximal to hadal environments in trenches adjacent to continental margins, providing a natural framework for examining microbial pressure responses across hydrostatic regimes and sediment burial depths.

Here, we experimentally assessed community-level responses to pressure across sediment depths ranging from the seafloor surface to 20 meters below the seafloor (mbsf) in the Puerto Rico Trench region (493–8,385 m water depth). Bioorthogonal non-canonical amino acid tagging (BONCAT) was used to compare the proportion of anabolically active cells in natural communities incubated without added complex organic growth substrates under atmospheric and *in situ* pressure. Stable-isotope tracing and nanoscale secondary ion mass spectrometry (nanoSIMS) provided complementary single-cell measurements in surficial trench-axis sediments. Fluorescence-activated cell sorting (FACS), metagenomics, and comparative phylogenomics were used to resolve the populations underlying these pressure responses. Together, these approaches reveal a reversal in microbial pressure responses with burial, from piezosensitivity at the sediment surface to piezophily in the subseafloor, and link this response to widespread hadal JS1/Atribacterota lineages.

## Results

To examine microbial pressure responses across an *in situ* hydrostatic pressure gradient, we investigated seven sites extending from the margin-proximal site PR02, across the bathyal slope (PR05 and PR04) and abyssal slope (PR06), to the hadal trench axis (PR07 and PR08) and adjacent abyssal plain (PR10) (Fig. 1a,b; Supplementary Table 1). These sites differed in organic matter content and composition, redox conditions, microbial biomass proxies and diversity, providing contrasting depositional and biogeochemical settings (Figs. 1c and 2).

**Fig. 1.**
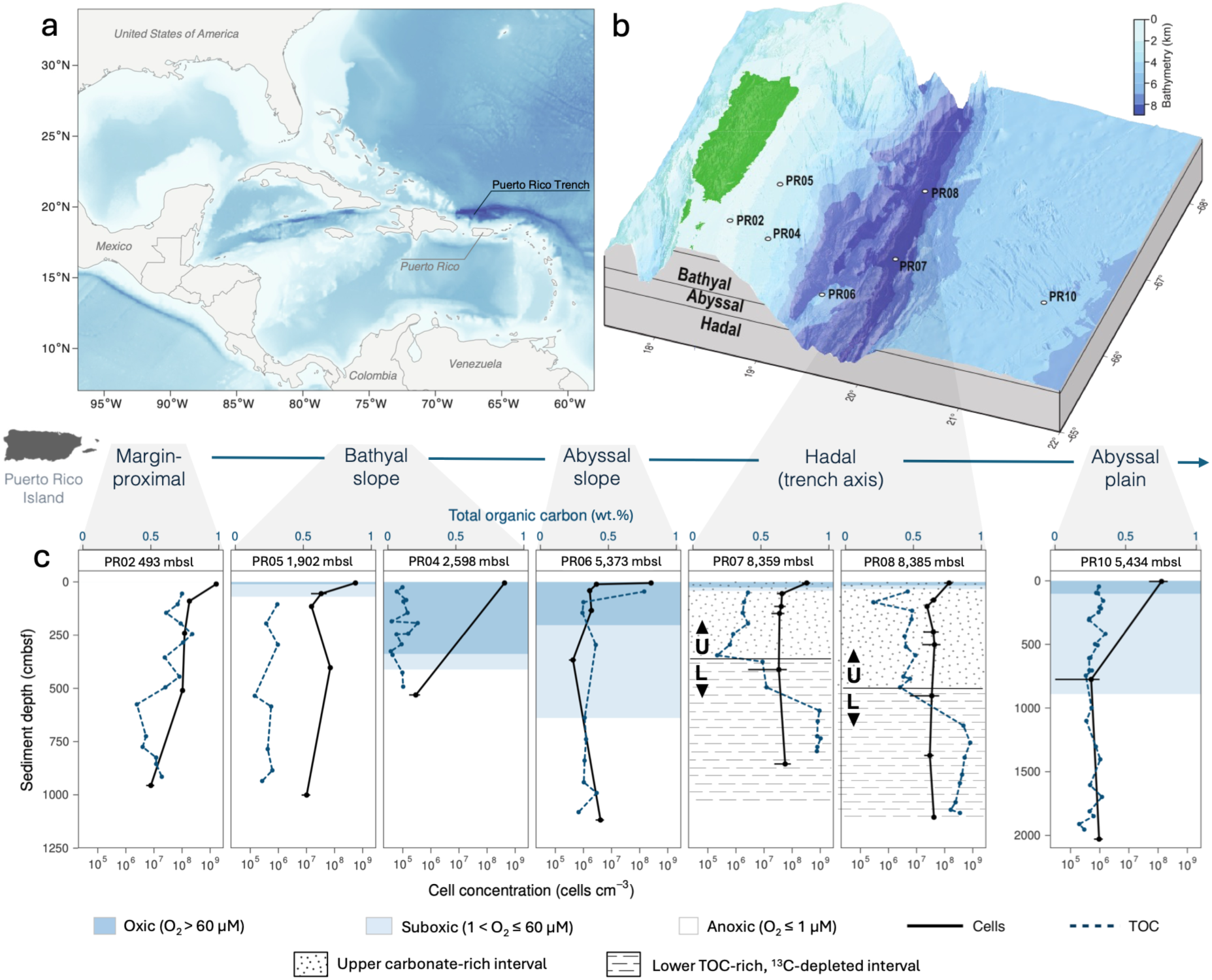
Organic matter, redox and microbial biomass vary across depositional settings, with pronounced heterogeneity at the trench axis. a, Regional location of the Puerto Rico Trench. b, Bathymetry and sampling sites. c, Sediment-depth profiles of cell concentration and total organic carbon content (TOC; wt.%) across redox zones and depositional intervals. Cell concentrations are shown as means of duplicate measurements; TOC content represents single measurements.

**Fig. 2.**
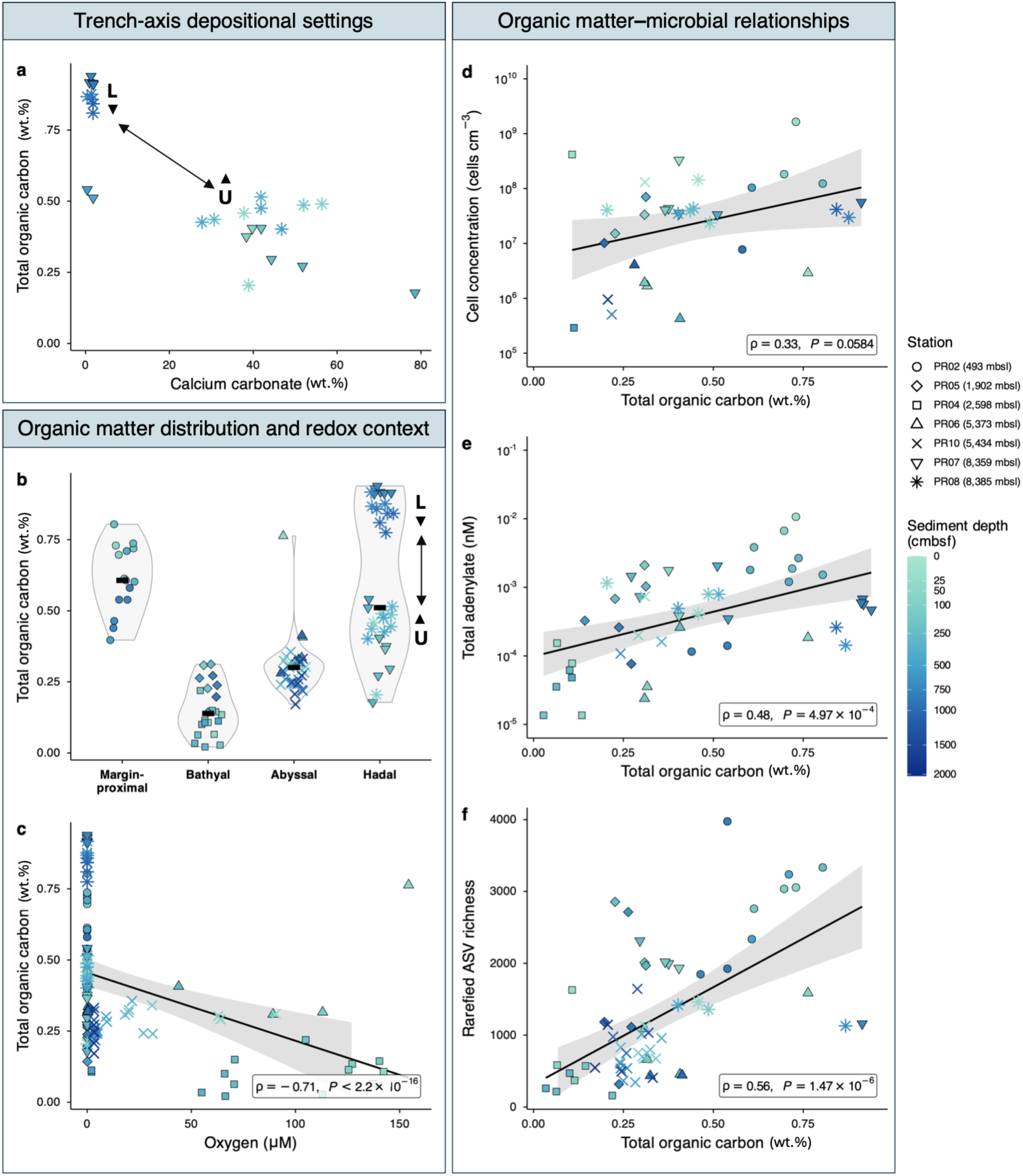
Organic matter covaries with microbial biomass and diversity across the transect but decouples in the lower trench-axis sediment interval. **a**, Relationship between total organic carbon (TOC; wt.%) and calcium carbonate (CaCO_3_; wt.%) in trench-axis sediments. **b**, TOC distributions across depositional settings. **c**, Relationship between TOC and oxygen concentration (µM). **d–f**, Relationships between TOC and cell concentration (cells cm^-3^), total adenylate (ATP + ADP + AMP; nM) and rarefied amplicon sequence variant (ASV) richness, respectively. Black lines indicate linear regressions, with shading representing 95% confidence intervals; insets report Spearman ρ and *P* values. Correlation analyses include only samples with paired measurements.

### Pressure effects on the proportion of anabolically active cells reverse with burial

We quantified the proportion of anabolically active cells using BONCAT in paired atmospheric- and *in situ*-pressure incubations of surficial and subsurface sediments (Figs. 3 and 4; Supplementary Table 8). Responses yielding higher proportions of anabolically active cells under atmospheric or *in situ* pressure were classified as piezosensitive or piezophilic, respectively, following the activity-based pressure-response framework of Amano et al.^5^. Across all incubations, BONCAT-positive cells comprised 0.4–56.8% of cells.

**Fig. 3.**
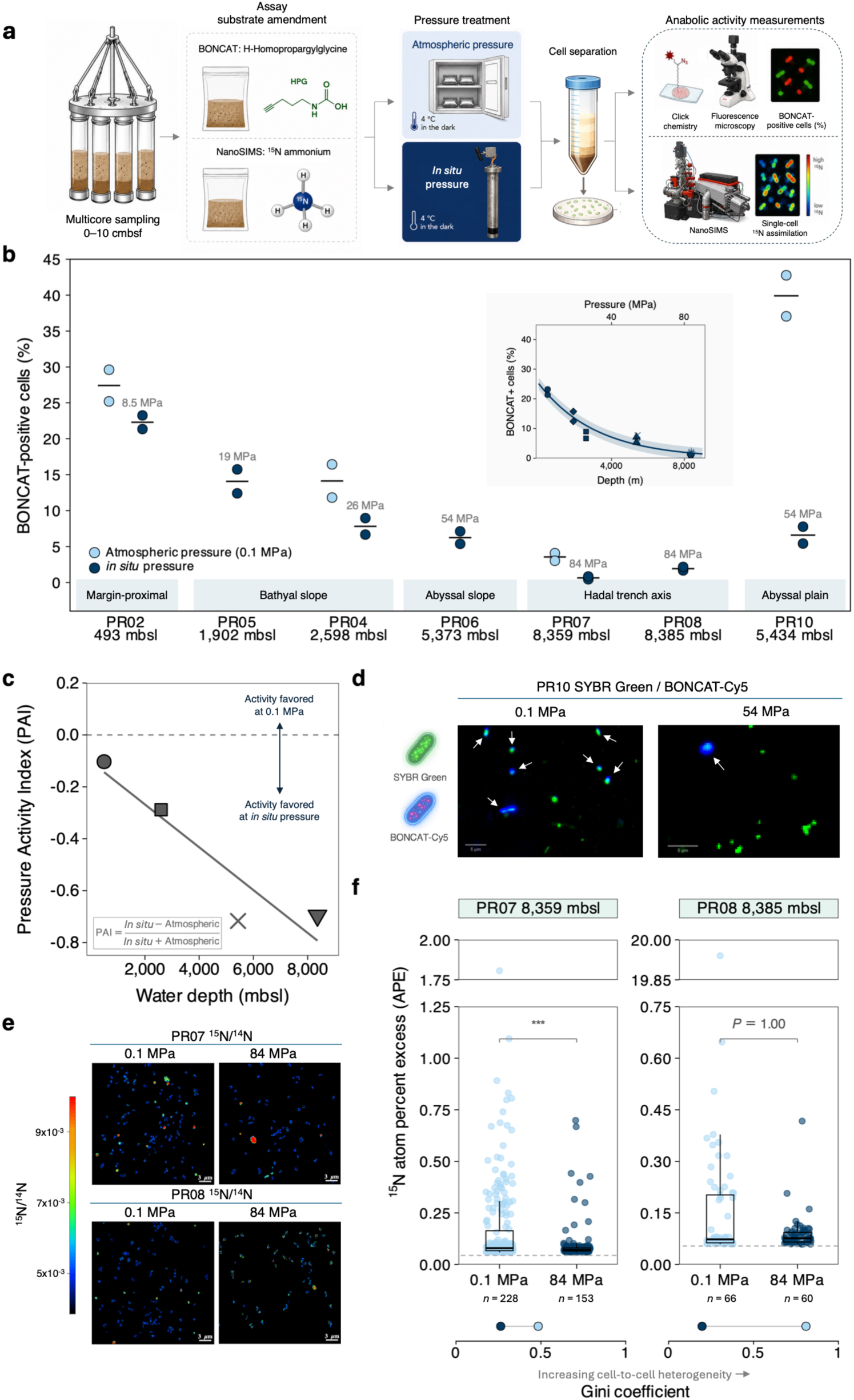
Atmospheric pressure increases anabolic activity in surficial sediments. **a**, Workflow for paired atmospheric and *in situ* pressure incubations of surficial sediments (0–10 cmbsf) followed by BONCAT and nanoSIMS. **b,** BONCAT-positive cells across the transect (two independent incubation replicates per sample and pressure treatment); inset shows the relationship between the *in situ*-pressure BONCAT-positive proportion and water depth. **c,** Pressure Activity Index (PAI), calculated from the mean proportion of BONCAT-positive cells for each pressure treatment; negative and positive values indicate greater activity under atmospheric and *in situ* pressure, respectively. **d,** Representative BONCAT fluorescence images from PR10; arrows indicate BONCAT-positive cells. **e**, Representative nanoSIMS ¹⁵N/¹⁴N images from PR07 and PR08; areas outside regions of interest (ROIs) interpreted as cells were set to zero and are shown in black. **f**, Cellular ¹⁵N atom percent excess (APE) among ¹⁵N-active ROIs from PR07 and PR08; dashed lines indicate the threshold used to classify ¹⁵N-active ROIs. Boxes show medians and interquartile ranges; *n* indicates ROIs analyzed. Each site and pressure represents one incubation; statistical comparisons are based on cell-level distributions rather than independent incubation replicates. Two-sided Wilcoxon rank-sum tests with Bonferroni correction (\*\*\**P* < 0.001; PR08, adjusted *P* = 1.00). Gini coefficients indicate cell-to-cell heterogeneity in ¹⁵N assimilation.

**Fig. 4.**
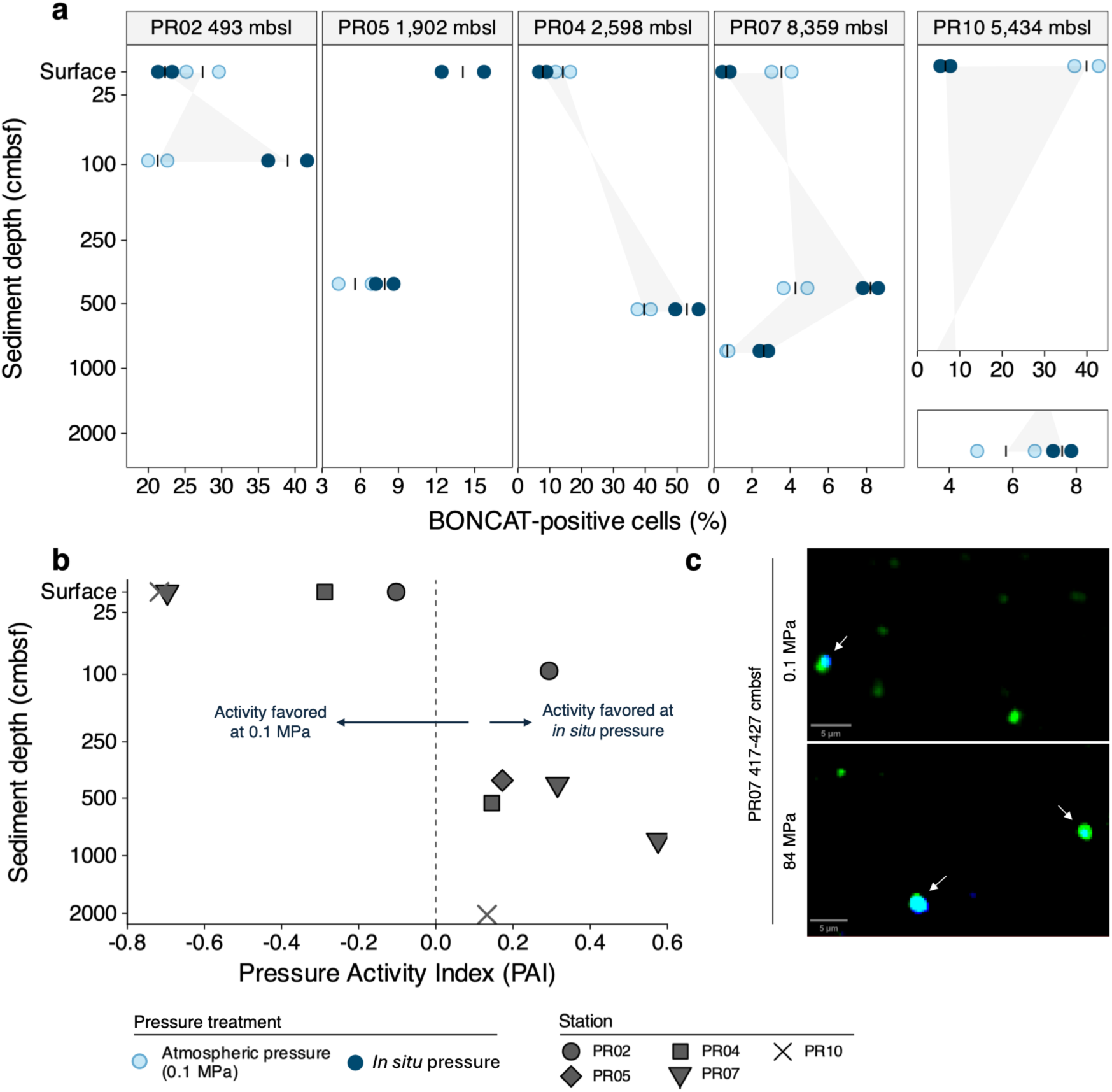
Microbial pressure responses reverse with burial from atmospheric-pressure-favored to *in situ*-pressure-favored activity. **a**, BONCAT-positive cells in surface and subsurface sediments under atmospheric and *in situ* pressure (two independent incubations per sample and pressure treatment). Grey shaded areas connect atmospheric- and *in situ*-pressure treatment means across sediment depths, highlighting the reversal in pressure response with burial. **b,** Pressure Activity Index (PAI) across sediment depths, calculated from the mean proportion of BONCAT-positive cells for each pressure treatment; negative and positive values indicate greater activity under atmospheric and *in situ* pressure, respectively. **c,** Representative BONCAT fluorescence images from PR07 at 417–427 cmbsf under atmospheric and *in situ* pressure; arrows indicate BONCAT-positive cells.

In surficial sediments, the proportion of BONCAT-positive cells under *in situ* pressure decreased with increasing water depth, from approximately 22% at the shallowest site (PR02; 493 meters below sea level [mbsl]) to <2% at the hadal trench-axis sites PR07 and PR08 (8,359–8,385 mbsl) (ρ = −0.93, *P* = 0.0025, *n* = 7 sites; Fig. 3b). Among the four sites with paired atmospheric and *in situ* pressure treatments (PR02, PR04, PR10 and PR07), mean proportions of BONCAT-positive cells were 1.2–6.1-fold higher under atmospheric pressure, with the greatest differences at sites PR10 and PR07. The Pressure Activity Index (PAI), calculated from mean BONCAT-positive proportions as (*in situ* − atmospheric)/(*in situ* + atmospheric), became increasingly negative across the *in situ* pressure gradient along the transect, reaching −0.72 at PR10 (54 MPa) and −0.70 at PR07 (84 MPa; Fig. 3c).

In contrast, all six subsurface horizons yielded 1.3–3.7-fold higher mean BONCAT-positive proportions under *in situ* pressure across margin-proximal, bathyal-slope, hadal trench-axis and abyssal-plain sediments (PR02, PR05, PR04, PR07 and PR10; Fig. 4a; Supplementary Table 8). PAI values were consistently positive (0.13–0.57), demonstrating a reversal in pressure response with burial (Fig. 4b).

### Atmospheric pressure enhances single-cell anabolic activity in surficial sediment at trench-axis PR07

We independently assessed surficial BONCAT-derived piezosensitivity using nanoSIMS measurements of single-cell ¹⁵N-ammonium and ¹³C-bicarbonate assimilation at PR07 and PR08. At PR07, median ¹⁵N atom percent excess (APE) among ¹⁵N-active cells was higher under atmospheric (0.0792) than *in situ* pressure (0.0710) in cell-level comparisons of single incubations (adjusted P = 4.13 × 10⁻⁵; Fig. 3f; Supplementary Table 9). At PR08, the cell-level distributions did not differ between pressure treatments (adjusted *P* = 1.00). Nevertheless, atmospheric-pressure incubations at both sites showed broader ¹⁵N APE distributions and greater cell-to-cell heterogeneity than *in situ*-pressure incubations, with extended upper tails particularly apparent at PR08 (Supplementary Table 9). In contrast, ¹³C-bicarbonate assimilation was negligible across pressure treatments at both sites (Supplementary Table 9).

### Active microbial populations differ between pressure regimes in trench-axis sediments

To identify populations underlying pressure responses at trench-axis PR07, BONCAT-positive cells from both pressure treatments were isolated and sequenced using fluorescence-activated cell sorting (BONCAT-FACS-seq). Libraries contained 211–3,078 sorted cells per fraction, and genera detected in the sequenced negative control were excluded (Supplementary Table 10). In surficial sediment, *Omnitrophus* dominated the *in situ*-pressure fraction (93.6% of Bracken-estimated genus-level reads), whereas *Pseudomonas* dominated under atmospheric pressure (79.3%), with a smaller contribution from *Psychrobacter*. In subsurface sediments (417–427 and 825–835 cmbsf), *in situ*-pressure fractions were dominated by JS1/Atribacterota, primarily the Genome Taxonomy Database (GTDB) placeholder taxon 34-128 (90–96%), whereas atmospheric-pressure fractions were dominated by *Acinetobacter* (82.8% and 99.9%, respectively), with smaller contributions from *Pseudomonas*. Previous studies reported atmospheric-pressure activity and piezosensitivity among hadal *Pseudomonas* and *Psychrobacter*^25,26^. However, *Pseudomonas* and *Acinetobacter* are potential low-biomass contaminants despite their absence from the reagent-only control, and secondary growth cannot be excluded^27,28^, precluding further physiological interpretation. In both subsurface horizons, *in situ*-pressure BONCAT-positive communities more closely resembled unincubated communities (T₀) than atmospheric-pressure communities, based on Bray–Curtis dissimilarity (Supplementary Table 10). Similarity to T₀ under *in situ* pressure was greater in the subsurface than at the surface.

To identify resident active populations, BONCAT-FACS-seq reads were recruited to the dereplicated T₀ metagenome-assembled genome (MAG) collection. No surficial BONCAT-FACS-seq reads, including those assigned to *Omnitrophus*, recruited to the available MAG collection, potentially reflecting incomplete genomic representation of active surficial populations. Prolonged incubations may also have introduced community selection alongside physiological responses. At both subsurface horizons, JS1 MAGs within the GTDB 34-128 lineage accounted for 53.9% and 64.0% of MAG relative abundance in *in situ*-pressure BONCAT-positive fractions at 417–427 and 825–835 cmbsf, respectively (Supplementary Table 11). Reads from below-threshold (B-T) fractions also recruited to these JS1 MAGs, indicating that these populations included cells both below and above the BONCAT-positive fluorescence threshold. High-confidence recruitment identified resident JS1 MAGs associated with anabolic activity under *in situ* pressure: PRT_8359_394_Bin36 and PRT_8359_709_Bin1 at 417–427 cmbsf, and PRT_8359_394_Bin36 at 825–835 cmbsf.

### Organic matter and redox structure vary across depositional settings

To contextualize pressure-dependent activity, we examined relationships among organic matter properties, redox conditions, and microbial biomass across the transect. Total organic carbon (TOC) content was generally higher at the margin-proximal site PR02 (0.40–0.80 wt.%), intermediate at PR05, PR06 and PR10, and lowest at PR04 (0.02–0.22 wt.%), whereas the highest individual TOC values occurred in deeper horizons at the hadal trench-axis sites PR07 and PR08 (up to 0.94 wt.%; Figs. 1c and 2b; Supplementary Table 2). Bulk organic carbon isotopic composition (δ¹³Corg) was relatively ¹³C-enriched at PR02 and PR05 (−23.3 to −19.0‰), generally intermediate at PR04, PR06 and PR10 (−28.2 to −22.4‰), with occasional strongly ¹³C-depleted values, and consistently more ¹³C-depleted in the lower TOC-rich intervals at the hadal trench-axis sites PR07 and PR08 (−27.7 to −23.5‰; Extended Data Fig. 1a; Supplementary Table 2).

Sites PR07 and PR08 exhibited an upper carbonate-rich interval and a lower TOC-rich, ¹³C-depleted interval separated at 425 and 575 centimeters below seafloor (cmbsf), respectively (Fig. 1c; Supplementary Table 2). The upper interval contained high CaCO₃ content (up to 78.6 wt.%), moderate TOC content (0.18–0.54 wt.%) and relatively ¹³C-enriched organic carbon (δ¹³Corg, −24.2 to −22.2‰), whereas the lower interval contained low CaCO₃ content (0.4–1.9 wt.%), higher TOC content (0.77–0.94 wt.%) and more ¹³C-depleted organic carbon (−27.7 to −27.1‰; Fig. 2a; Extended Data Fig. 2; Supplementary Table 2).

Across the full dataset, TOC content was strongly negatively correlated with oxygen concentration (ρ = −0.71, *P* < 2.2 × 10⁻¹⁶; Fig. 2c; Supplementary Table 4), whereas δ¹³C_org_ showed a weaker negative correlation (ρ = −0.35, *P* = 2.37 × 10⁻⁴; Supplementary Table 4). Oxygen penetration was shallower at PR02, PR05, PR07 and PR08 than at PR04, PR06 and PR10 (Extended Data Fig. 1c; Supplementary Table 3). It broadly decreased with increasing TOC content but differed among intermediate-TOC sites, being shallower at PR05 than at the more ¹³C-depleted PR06 and PR10.

### Biomass and diversity covary with organic matter properties

Cell abundance, total adenylate (ATP + ADP + AMP), and rarefied amplicon sequence variant (ASV) richness generally remained elevated downcore at sites PR02, PR05, PR07 and PR08 relative to PR04, PR06 and PR10 (Fig. 1c; Extended Data Fig. 1e,g; Supplementary Tables 5–7). Across the transect, TOC correlated positively with total adenylate (ρ = 0.48, *P* = 4.97 × 10⁻⁴) and rarefied ASV richness (ρ = 0.56, *P* = 1.47 × 10⁻⁶), but its relationship with cell abundance was weaker and not statistically significant (ρ = 0.33, *P* = 0.0584; Fig. 2d–f). δ¹³Corg correlated positively with cell abundance (ρ = 0.48, *P* = 0.014), total adenylate (ρ = 0.48, *P* = 8.3 × 10⁻⁴) and rarefied ASV richness (ρ = 0.56, *P* = 6.8 × 10⁻⁸; Extended Data Fig. 3j–l; Supplementary Table 4). Thus, TOC captured broad variation in microbial biomass-linked and diversity proxies, while δ¹³C_org_ tracked contrasting biological and redox conditions among sites with similar TOC.

### TOC content decouples from microbial biomass and richness in subseafloor ¹³C-depleted trench sediments

Despite higher TOC content, cell abundance, total adenylate, and ASV richness in the lower TOC-rich, ¹³C-depleted intervals at PR07 and PR08 fell below their transect-wide relationships with TOC (Fig. 2d–f). Averaged across sediment samples from PR07 and PR08 within each interval, total adenylate concentration decreased from 9.44 × 10⁻⁴ to 4.24 × 10⁻⁴ nM, and rarefied ASV richness decreased from 1,571 to 1,432 ASVs from the upper carbonate-rich to the lower TOC-rich, ¹³C-depleted intervals. In contrast, subsurface cell abundance remained relatively stable across this transition (Extended Data Fig. 1e; Supplementary Table 5). Adenylate content normalized to cell abundance also decreased across this transition (Extended Data Fig. 1f; Supplementary Table 7).

### Distinct JS1 populations are enriched in hadal trench-axis sediments

16S rRNA gene profiles and MAG distributions revealed differences in JS1/Atribacterota abundance and clade composition among PR05, PR07 and PR08 (Fig. 5d; Supplementary Table 12). Three dominant 99%-clustered JS1-16S clades (1–3) showed clear bathyal– hadal partitioning. At PR05, JS1-16S clade 3 predominated and total JS1 abundance remained low (0–14.11% of 16S reads), whereas trench-axis PR07 and PR08 were dominated by JS1-16S clades 1 and 2, reaching downcore maxima of 77.03% and 76.63%, respectively. This bathyal–hadal pattern was reflected among four representative Puerto Rico Trench JS1 MAGs: the two *in situ*-pressure-active MAGs at PR07 were present at low relative abundance at PR05 but became increasingly abundant downcore at PR07 and PR08 (Fig. 6c).

**Fig. 5.**
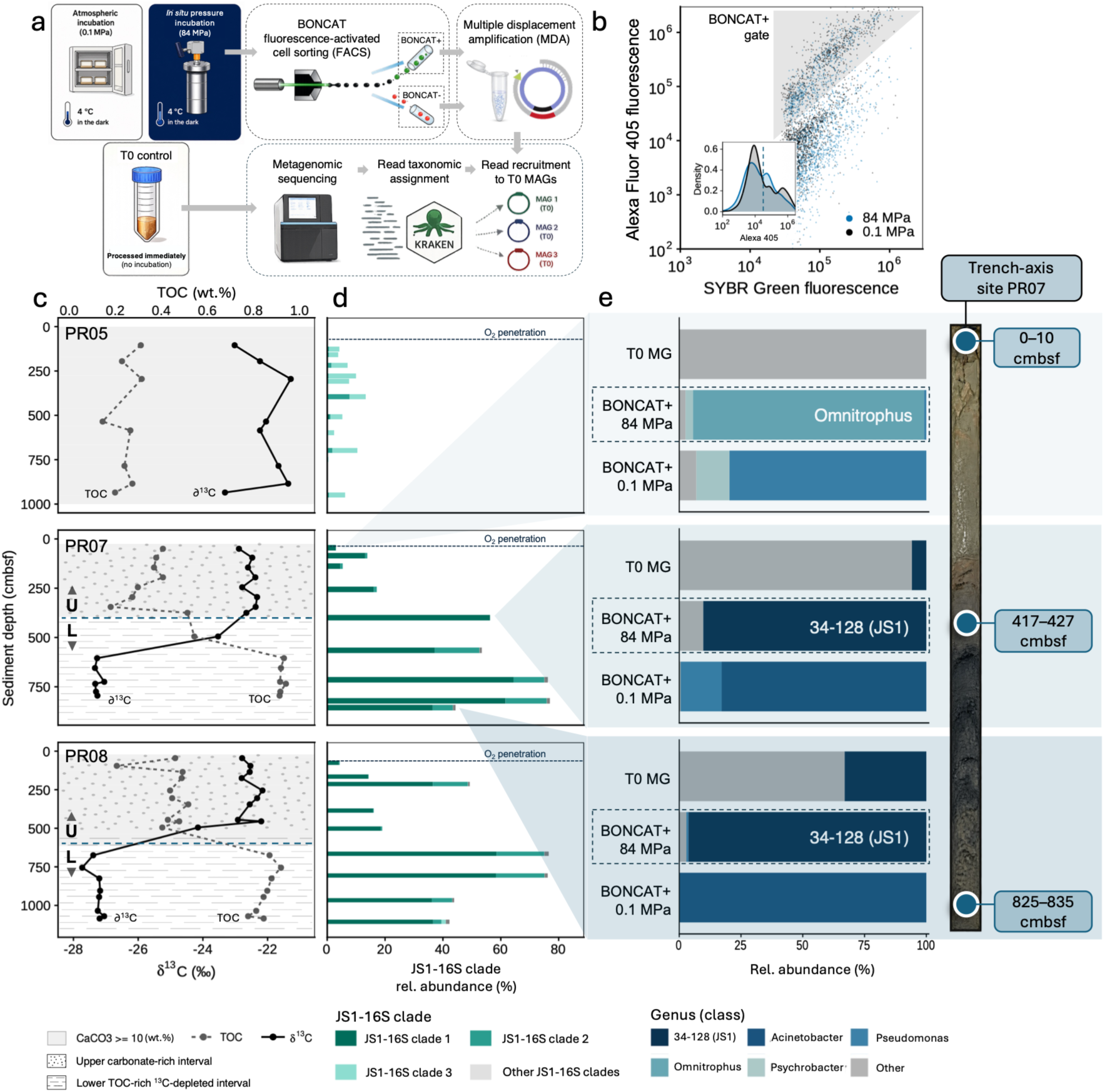
Abundant trench-axis JS1 populations are associated with distinct depositional intervals and anabolic activity under *in situ* pressure. **a**, BONCAT-FACS-seq workflow for atmospheric and *in situ* pressure incubations and T₀ metagenomic controls. **b**, Representative BONCAT-positive FACS gate and Alexa Fluor 405 fluorescence distributions. **c**, Total organic carbon (TOC; wt.%) and δ¹³C_org_ (‰) profiles across depositional intervals at PR05, PR07 and PR08. **d**, Relative abundance of JS1 16S rRNA gene clades with sediment depth. **e**, Genus-level composition of T₀ metagenomes and BONCAT-positive fractions under atmospheric and *in situ* pressure at trench-axis site PR07.

**Fig. 6.**
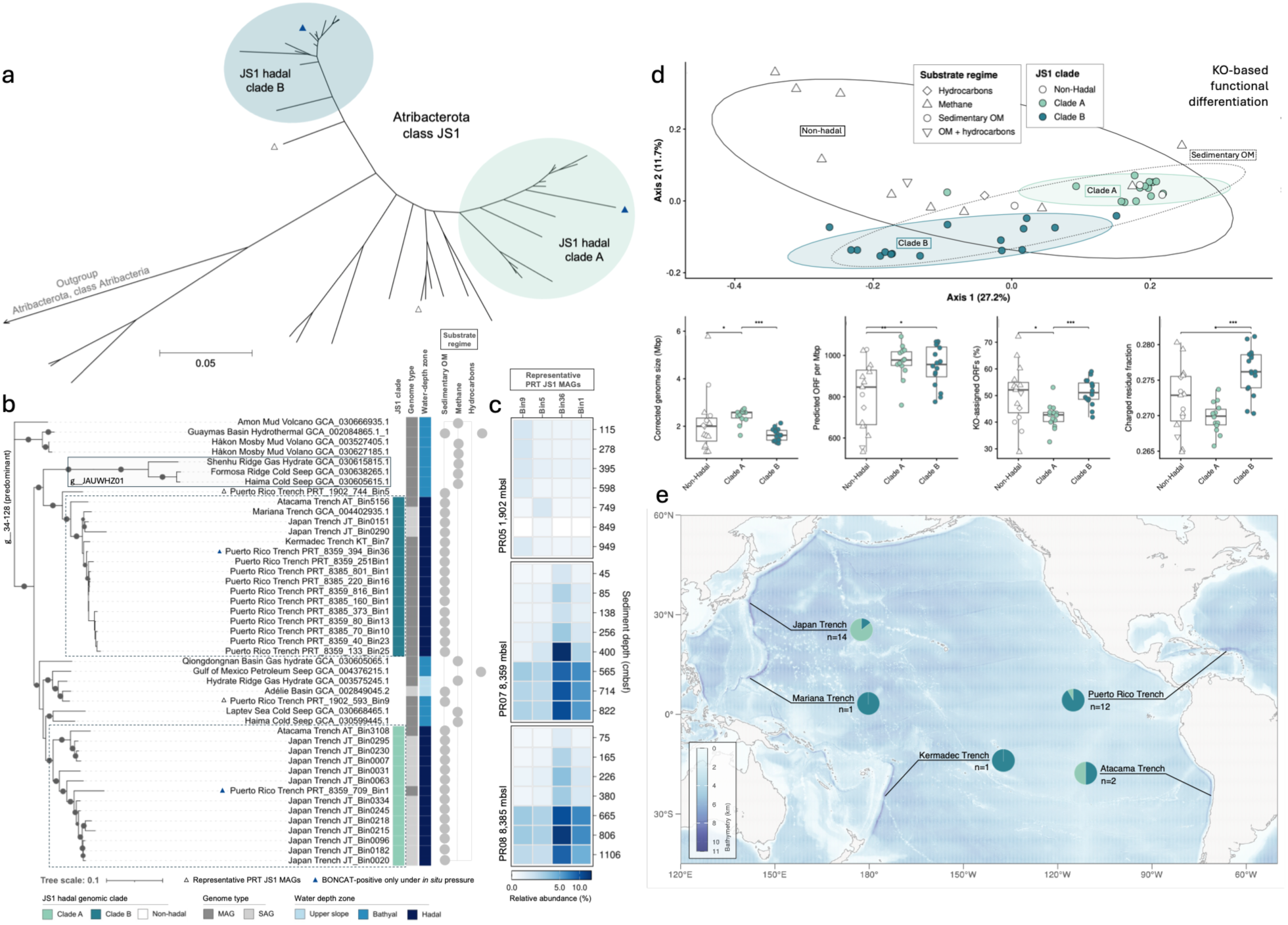
*In situ*-pressure-active JS1 populations belong to genomically differentiated clades widespread across hadal trenches. **a**, Simplified phylogenomic topology of JS1, highlighting hadal clades A and B. **b,** Detailed maximum-likelihood JS1 phylogeny annotated by JS1 clade, genome type, water-depth zone and substrate regime. Node size represents ultrafast bootstrap support based on 1,000 replicates. Filled blue triangles indicate Puerto Rico Trench MAGs detected in BONCAT-positive fractions under *in situ* but not atmospheric pressure. **c,** Relative abundance of representative PRT JS1 MAGs across sediment depths at PR05, PR07 and PR08. **d,** KEGG Ortholog (KO)-based functional differentiation and genome properties among non-hadal JS1 (*n* = 15 genomes), hadal clade A (*n* = 14) and hadal clade B (*n* = 16). The principal coordinates analysis (PCoA) shows Bray–Curtis dissimilarities of KO profiles. KO-profile differences were assessed by PERMANOVA; boxes show medians and interquartile ranges, with asterisks indicating pairwise Wilcoxon rank-sum tests with Benjamini–Hochberg-adjusted *P* values (\**P* < 0.05, \*\**P* < 0.01, \*\*\**P* < 0.001). **e,** Geographic distribution and number of publicly available JS1 MAGs assigned to hadal clades A and B across five hadal trenches.

JS1 abundance was highest within the lower TOC-rich, ¹³C-depleted intervals at PR07 and PR08 (Fig. 5c,d). Across JS1-containing sites (PR05, PR07 and PR08), JS1 relative abundance increased with TOC (ρ = 0.76, adjusted *P* < 0.001) and sediment depth (ρ = 0.53, adjusted *P* < 0.001), and decreased with oxygen (ρ = −0.77, adjusted *P* < 0.001) and δ¹³C_org_ (ρ = −0.56, adjusted *P* = 0.002; Supplementary Table 13).

### In situ-pressure-active JS1 belong to hadal-associated phylogenomic clades

Phylogenomic analysis of Puerto Rico Trench and geographically distant JS1 genomes resolved two hadal-associated clades spanning the Puerto Rico, Atacama, Japan, Kermadec and Mariana trenches (Fig. 6b,e; Supplementary Table 14). These lineages, designated JS1 hadal clades A and B, belonged to GTDB taxon 34-128 (Atribacterota, class JS1) and were distinct from non-hadal upper-slope and bathyal JS1 genomes. Pairwise average amino acid identity (AAI) comparisons placed most JS1 genome pairs, including comparisons between hadal clades A and B, within the genus-like range, whereas species-like values were restricted to subsets within each clade (Extended Data Fig. 6).

The two JS1 MAGs to which reads from *in situ*-pressure BONCAT-positive fractions were recruited belonged to the hadal-associated clades: PRT_8359_709_Bin1 in hadal clade A and PRT_8359_394_Bin36 in hadal clade B (Fig. 6b). In contrast, two representative JS1 MAGs recovered from the shallower PR05 site, PRT_1902_744_Bin5 and PRT_1902_593_Bin9, fell outside the hadal-associated clades and did not recruit reads from *in situ*-pressure BONCAT-positive fractions (Fig. 6b; Supplementary Table 11).

### Hadal-associated JS1 clades are genomically differentiated

KEGG Ortholog (KO)-based functional profiles differentiated JS1 phylogenomic groups, with hadal clade B showing the strongest separation from non-hadal genomes and clade A occupying a region closer to sedimentary organic-matter-associated non-hadal JS1 (Fig. 6d). KO-based functional profiles differed among JS1 genomic groups even after accounting for the carbon/substrate regime of their source environments (*e.g.,* methane-versus organic matter-dominated; PERMANOVA, genomic group R² = 0.156, P = 0.0001), although multivariate dispersion also differed among groups (P = 0.0003; Fig. 6d; Supplementary Table 15).

Clade B had the highest median predicted proteome-wide charged-residue fraction (27.62%), significantly exceeding both clade A (26.99%) and non-hadal genomes (27.29%), and these differences remained significant after accounting for genome completeness, contamination and genome recovery type (Fig. 6d; Supplementary Tables 16 and 17). Clade B also had a significantly higher KO-assigned open reading frame (ORF) fraction than clade A after accounting for these factors (Supplementary Table 17).

## Discussion

Here we report a reversal in microbial responses to hydrostatic pressure with burial across the Puerto Rico Trench region, from predominantly piezosensitive surficial communities to piezophilic subsurface communities. This reversal occurred across contrasting depositional settings, revealing piezophily as a recurring physiological feature of the subseafloor communities examined here. In subsurface hadal trench-axis sediments, dominant JS1/Atribacterota populations belonging to geographically widespread hadal lineages showed a strong response to *in situ* pressure, indicative of piezophily.

In surficial sediment at trench-axis site PR07, a community-level piezosensitive response was supported by both a higher proportion of anabolically active cells and greater ammonium assimilation among ¹⁵N-active cells under atmospheric relative to *in situ* pressure (Fig. 3b,c,f). Negligible ¹³C-bicarbonate assimilation across pressure treatments was consistent with heterotrophy predominating during the incubation period (Fig. 3f). This pattern mirrors early observations from surficial Puerto Rico Trench-axis sediments, where heterotrophic uptake and cellular assimilation of ^14^C-labeled amino acids were strongly suppressed at *in situ* relative to atmospheric pressure^16^. At the community level, such net pressure responses reflect the relative abundance and activity of pressure-sensitive and pressure-adapted populations^9^. Accordingly, continual delivery of particle-associated microorganisms with limited prior pressure exposure may sustain pressure-sensitive populations in surficial sediments across the transect, potentially enhanced by topographic focusing at the trench axis^15,22–24,29,30^. Consistent with this interpretation, the magnitude of surficial PAI increased with water depth, paralleling evidence from deep-water communities that increasing hydrostatic pressure progressively constrains microbial activity, with implications for estimates of carbon cycling at depth^5^. At PR07, greater cell-to-cell heterogeneity among ¹⁵N-active cells under atmospheric pressure further suggests that a highly responsive, pressure-sensitive subset may contribute disproportionately to the net pressure response (Fig. 3f)^31^.

Below the sediment surface, PAI captured the opposite pattern: *in situ* pressure yielded higher proportions of anabolically active cells across all six subsurface horizons examined (Figs. 4a,b). The consistent shift from piezosensitive surface responses to piezophilic responses after burial suggests that piezophilic physiology becomes advantageous in the subsurface, favoring the persistence of pressure-adapted members of subseafloor communities^9,12^. Sustained hydrostatic pressure over thousands to millions of years may filter surface-derived populations during burial, consistent with observed subseafloor community restructuring^32–34^. The present experiments cannot resolve whether pressure-adapted populations primarily persisted after burial or proliferated within the subsurface. However, the placement of pressure-active JS1 populations within geographically widespread hadal-associated phylogenomic clades links the observed physiological response to evolutionary differentiation among JS1 lineages. Together, these observations are consistent with sustained hydrostatic pressure acting as an ecological filter during burial.

Hydrostatic pressure also structured the taxonomic composition of the active fraction. At trench-axis site PR07, *Omnitrophus* dominated the *in situ*-pressure-active fraction in surficial sediment, while JS1/34-128 dominated the *in situ*-pressure-active fractions in both subsurface horizons. Neither was detected in the corresponding atmospheric-pressure-active fractions, consistent with piezophilic physiology in these populations (Fig. 5e; Extended Data Fig. 4d; Supplementary Tables 10 and 11). Active assemblages more closely resembled unincubated T₀ communities under atmospheric pressure at the surface and *in situ* pressure in the subsurface, suggesting that piezophilic physiology is embedded within the resident subseafloor community (Supplementary Table 10).

Recruitment of BONCAT-positive reads to resident T₀ MAGs further linked the subsurface pressure response to JS1/Atribacterota. Within this group, the piezophilic response was lineage-specific: genome-resolved populations active under *in situ* pressure at hadal PR07 were distinct from those characteristic of bathyal PR05. JS1-16S clades showed similar bathyal–hadal partitioning, consistent with ecological differentiation along the bathymetric gradient (Figs. 5d and 6b,c; Supplementary Tables 11 and 12). The pressure-active trench-axis populations belonged to hadal-associated phylogenomic clades spanning geographically distant trenches, raising the possibility that piezophilic physiology is widespread among hadal JS1 populations (Fig. 6b,e; Supplementary Table 14)^35–37^. Functional differentiation among JS1 genomic groups, strongest in hadal clade B, persisted after accounting for carbon/substrate regime, suggesting that hydrostatic pressure may contribute alongside other hadal environmental factors (Fig. 6d; Supplementary Table 15). Candidate traits for maintaining cellular function under pressure included an elevated charged-residue fraction in clade B and the genomic potential of Puerto Rico Trench JS1 MAGs to synthesize the compatible solutes trehalose and di-myo-inositol-phosphate, as identified in a separate study^38^, although their contributions to piezophily remain untested (Extended Data Fig. 7; Supplementary Tables 16 and 17)^39–41^. Previous studies have shown that JS1/Atribacterota are abundant and transcriptionally active in deeply buried marine sediments, including evidence for populations persisting over geological timescales^42,43^. Our findings provide evidence of piezophilic physiology among particular JS1 populations belonging to evolutionarily differentiated hadal lineages spanning geographically distant trenches, potentially contributing to their ecological success in the subseafloor biosphere.

The exceptional enrichment of both hadal JS1 clades at the trench axis coincided with sustained high pressure and distinctive depositional and energetic conditions. A carbonate-rich upper interval overlay a TOC-rich lower interval with δ¹³Corg reaching −27.7‰, the most depleted value among surveyed hadal sediments (Fig. 5c; Extended Data Fig. 1b; Supplementary Table 18). This succession may reflect episodic downslope delivery of margin- or terrestrial-derived organic matter, consistent with a previously identified regional debrite^22,23,44,45^. Despite high TOC, the lower interval supported disproportionately low microbial standing stocks, adenylate content and diversity, potentially reflecting organic matter source and lability alongside low anaerobic energy yields (Figs. 1c and 2d–f; Extended Data Fig. 1e–g; Supplementary Tables 5–7)^44,46–48^. Yet JS1/Atribacterota comprised up to 77% of 16S reads, among the highest abundances reported for marine sediments (Fig. 5d; Supplementary Tables 12 and 13)^35,40,49–51^. This enrichment despite low microbial standing stocks suggests that JS1 populations may be particularly well suited to persistence under these restrictive conditions. JS1 MAGs encoded pathways for peptide and sugar fermentation and microcompartment-associated secondary fermentation, potentially supporting energy conservation and persistence under these conditions^35,40,42,51,52^. The recurrence and ecological prominence of hadal JS1 lineages across geographically distant trenches may therefore reflect ecological specialization to both sustained hydrostatic pressure and trench-specific depositional, redox, and energetic regimes^53^.

Together, our findings identify hydrostatic pressure as a key determinant of microbial activity in marine sediments. Its effect reversed with burial, from piezosensitive responses at the sediment surface to piezophilic responses across contrasting subsurface settings, including sediments buried meters beneath the hadal trench axis. This reversal, together with the association of pressure-active JS1 populations with widespread hadal lineages, is consistent with the hypothesis that sustained pressure acts as an ecological filter over long timescales, favoring populations with piezophilic physiology. Because pressure altered the prevalence and taxonomic distribution of anabolic activity, *in situ*-pressure experiments are essential for constraining microbial contributions to sedimentary biogeochemical cycling. More broadly, our findings suggest that piezophilic physiology may be widespread in the deep subseafloor biosphere, providing a new perspective on microbial adaptation and persistence in one of Earth’s largest microbial habitats.

## Methods

### Study area and sample collection

Sediment samples were collected from seven sites (PR02, PR04, PR05, PR06, PR07, PR08 and PR10) spanning margin, slope, hadal trench-axis and adjacent abyssal-plain settings of the Puerto Rico Trench region during cruise AR64-02 (15 February–10 March 2022) aboard the R/V *Neil Armstrong* (Fig. 1a,b; Supplementary Table 1). Sampling sites ranged from 493 to 8,385 meters below sea level (mbsl), corresponding to approximate *in situ* hydrostatic pressures of 5–84 MPa. Sediment cores were recovered using multicore, gravity-core and piston-core systems.

Recovered cores were sectioned into whole rounds on deck and maintained at 4 °C in a shipboard refrigerated van until subsampling. Core temperature was monitored upon recovery to minimize warming during retrieval and handling. Sediment subsamples were collected from the interior of each core section using sterile 60-mL cut-off syringes to minimize contamination from material in contact with the core liner and were subsequently processed or preserved according to downstream analyses.

### Ancillary geochemical, biochemical, and molecular datasets

Geochemical, biochemical, and molecular datasets used in this study included dissolved oxygen (O₂), nitrate (NO₃⁻), sulfate (SO₄²⁻), TOC, total nitrogen (TN), CaCO₃, δ¹³C_org_, total adenylate (ATP + ADP + AMP), 16S rRNA gene profiles, and sediment metagenomes. O₂, NO₃⁻, and SO₄²⁻ concentrations were measured as previously described by Fischer et al.^54^. Total organic carbon (TOC) content and δ¹³Corg were determined following the method described in Swenson Perger et al.^55^ and Cohen et al.^56^ in the Department of Geosciences at Stony Brook University. TN and CaCO_3_ were measured as described by Trejos-Espeleta et al.^57^. Total adenylate was measured as described by Orsi^58^ and normalized to cell abundance where paired measurements were available. Additionally, 16S rRNA gene profiles and the Puerto Rico Trench MAG collection generated from field-collected, unincubated sediment samples were incorporated from Breusing et al.^38^; metagenomic assembly, genome binning, quality assessment, and dereplication are described therein.

### 16S rRNA gene analysis and JS1 clade definition

Previously generated 16S rRNA gene profiles from the broader Puerto Rico Trench amplicon dataset (NCBI BioProject PRJNA1334354) were reanalyzed to evaluate depth distributions of JS1-affiliated populations across PR05, PR07 and PR08^38^. JS1-affiliated ASVs were extracted from the finalized ASV table, and per-sample relative abundances were calculated from read counts normalized to total reads. To evaluate fine-scale JS1 population structure, JS1-affiliated sequence variants were clustered by pairwise sequence similarity using VSEARCH at a 99% identity threshold^59^. The three dominant 99%-identity groups are referred to here as JS1-16S clades 1–3 to distinguish them from genome-based hadal clades A and B.

### Sediment cell separation

Microbial cells were separated from sediment using a workflow modified from Morono et al.^60^. The workflow was used for total cell enumeration of unincubated samples and downstream BONCAT and nanoSIMS analyses of incubated samples. Cell separation for total counts was performed at JAMSTEC, whereas BONCAT-associated extractions were performed at the Scripps Institution of Oceanography (SIO). Formaldehyde-fixed sediment slurry was combined with a chemical cell-detachment mixture containing 100 mM EDTA, 100 mM sodium pyrophosphate, 1% (v/v) Tween 80 and methanol and vortexed at high speed for 1 h. Samples were subsequently sonicated for 20 min in 30-s on/off cycles at 4 °C. A detailed protocol is available on protocols.io^61^. Samples processed at JAMSTEC were sonicated in an ice-cooled Bioruptor water bath (Diagenode, Seraing, Belgium) at 20 W, whereas samples processed at SIO were sonicated on ice using a Model 50 Sonic Dismembrator (Fisher Scientific, Waltham, MA, USA) at 20% power. Cell extracts were separated by density-gradient centrifugation over 30%, 50% and 80% Nycodenz layered above 80% (w/v) sodium polytungstate at 10,000 × *g* for 1 h at 4 °C. The recovered aqueous cell fraction was used for downstream analyses.

### Total cell enumeration

For total cell counts, the recovered cell fraction was diluted 1:100, collected onto 0.2-µm polycarbonate filters, stained with 10× SYBR Green I in phosphate-buffered saline (PBS), and enumerated at JAMSTEC using an automated fluorescence microscopy system based on an Olympus BX-51 microscope (Olympus Corporation, Hachioji, Tokyo, Japan) equipped with an automatic slide handler, LED illumination and a cooled CCD camera, following Morono et al. and Morono and Inagaki^60,62^.

### Pressure incubations

Before the cruise, artificial seawater (ASW) was prepared from Sea Salts (40 g L⁻¹; Sigma-Aldrich, St. Louis, MO, USA; pH ∼7), amended with L-cysteine hydrochloride (0.5 g L⁻¹) as a reducing agent and resazurin (0.5 mg L⁻¹) as an oxygen indicator, aliquoted in 30-mL volumes into metalized polyethylene terephthalate pouches (IMPAK Corporation, Sebastian, FL, USA), flushed with ultra-high-purity N₂ and heat-sealed. The pre-aliquoted ASW was kept cold until sediment processing aboard ship. Pouches were then reopened under continuous N₂ flushing, and sediment slurry incubations were prepared by combining 30 mL of sediment with 30 mL of ASW (1:1, v/v). Sodium sulfide (150 µL of a 5% [w/v] stock solution; Sigma-Aldrich) was added as a reductant to maintain anoxic conditions. BONCAT incubations were amended with L-homopropargylglycine (HPG; Vector Laboratories, Newark, CA, USA; CCT-1067) to a final concentration of 50 µM, whereas stable-isotope incubations for nanoSIMS were amended with ¹³C-bicarbonate (1.04 mM; Cambridge Isotope Laboratories, Tewksbury, MA, USA; CLM-441) and ¹⁵N-ammonium (0.65 mM; Cambridge Isotope Laboratories; NLM-467). All shipboard manipulations were performed on ice using pre-chilled ASW, with sediment slurries continuously flushed with ultra-high-purity N₂ during processing to minimize oxygen exposure. Following addition of sediment and incubation amendments, pouches were homogenized and heat-sealed without headspace. An ASW-only control was included with each pressure-treatment set to assess maintenance of anoxic conditions independently of the sediment incubation controls.

*In situ*-pressure incubations were conducted in custom pin-retained piston-closure pressure vessels filled with distilled water as the hydraulic medium^63^. Sealed incubation pouches were loaded into the vessels and pressurized to *in situ* hydrostatic pressure using an eTensifier electric high-pressure pump system (High Pressure Equipment Company [HiP], Erie, PA, USA). For the shallowest site (PR02; 493 m water depth), the surface sediment incubation was conducted at 8.5 MPa, the minimum pressure achievable with the pressure intensifier, rather than the approximately 5 MPa corresponding to the sampling depth. Pressure was measured using the eTensifier integrated pressure transducer. For BONCAT, each sample included two HPG-amended replicates and one HPG-free sediment negative control at *in situ* pressure, with a parallel set maintained at atmospheric pressure. Both atmospheric pressure and *in situ* pressure incubations were maintained at 4 °C for the remainder of the cruise. Pressurized vessels were transported under pressure in a refrigerated van to Scripps Institution of Oceanography, where incubation continued at 4 °C for a total incubation period of 10 weeks to permit detectable HPG incorporation in low-activity deep-sediment communities. Vessel pressure was verified using the eTensifier immediately before depressurization and subsampling at the end of the incubation period. ASW-only controls were opened in an anaerobic chamber and inspected for resazurin color change to verify maintenance of anoxic conditions.

After incubation, pouches were opened in an anaerobic chamber (Coy Laboratory Products, Grass Lake, MI, USA), and matching 5-mL slurry aliquots were collected for downstream analyses. Aliquots for BONCAT and nanoSIMS activity measurements were fixed overnight at 4 °C with formaldehyde at a final concentration of 4%, washed with PBS and 1:1 PBS:ethanol, and preserved in ethanol. Unfixed aliquots for BONCAT-FACS-seq were preserved separately in glycine-TE buffer.

### BONCAT single-cell activity measurements

Extracted cells were prestained with SYBR Green I in solution, collected onto 25-mm, 0.2-µm polycarbonate filters, and washed twice with 3 mL of 1× PBS before on-filter click chemistry following the general framework of Couradeau et al.^64^; a detailed protocol is available at protocols.io^65^. The click-reaction mixture contained 100 µM CuSO₄, 500 µM THPTA, 5 µM Cy5 azide dye (Vector Laboratories, Newark, CA, USA), 5 mM sodium ascorbate, and 5 mM aminoguanidine-HCl in 1× PBS. Filters were incubated with 80 µL of click-reaction mixture for 30 min in the dark under a coverslip. After labeling, filters were washed three times with 3 mL of 1× PBS, cut into quarters, and one quarter was mounted on glass slides in VECTASHIELD mounting medium; the remaining filter material was preserved at −80 °C.

Mounted filter quarters were imaged on a SLIDEVIEW VS200 fluorescence microscope (Olympus Corporation, Hachioji, Japan) using FITC and Cy5 filter sets: FITC/SYBR Green, Ex 474/27 nm, dichroic 495 nm, Em 525/45 nm; and Cy5, Ex 635/18 nm, dichroic 652 nm, Em 680/42 nm. For each replicate filter, a large-area scan covering most of one mounted quarter was acquired and analyzed in QuPath^66^. Total cells were identified from the SYBR Green/FITC channel using the Cell Detection tool, and BONCAT-positive cells were identified from Cy5 fluorescence using a single-image object classifier.

Corresponding HPG-free negative controls were carried through labeling and imaging and used to define background Cy5 fluorescence and BONCAT-positive classification thresholds, which were held constant across replicate images within each sample and treatment. Counts of total and BONCAT-positive cells were used to calculate the proportion of anabolically active cells. Pressure effects were summarized using a normalized Pressure Activity Index (PAI), adapted from the pressure-effect metric (Pe) of Tamburini et al.^9^. PAI was calculated from the mean proportion of BONCAT-positive cells across the two incubation replicates for each treatment as (*in situ* − atmospheric)/(*in situ* + atmospheric), with positive and negative values indicating greater activity under *in situ* and atmospheric pressure, respectively.

### NanoSIMS single-cell activity measurements

Fixed cells collected on gold-coated filters were mounted on conductive carbon tabs on glass slides and analyzed using a NanoSIMS 50L (CAMECA, Gennevilliers, France) at the nano@stanford facility. Randomly selected areas were pre-sputtered over 40 × 40 µm for 2 min, after which 30 × 30 µm areas were analyzed with a 5-pA Cs⁺ primary ion beam, a 200-µm aperture (D1-3), and a 256 × 256-pixel raster across 40 serial images. An entrance slit of 20 × 140 µm (ES-3) and aperture slit of 80 × 80 µm (AS-4) were used. Secondary ions were collected for ¹²C⁻, ¹³C⁻, ¹²C¹²C⁻, ¹²C¹³C⁻, ¹²C¹⁴N⁻ and ¹²C¹⁵N⁻. Mass resolving power was approximately 9,000 as reported by the CAMECA software. Regions of interest (ROIs) that were interpreted as cells were manually delineated in Look@nanoSIMS^67^ using carbon-, nitrogen- and isotope-ratio images. Between 194 and 377 ROIs were analyzed per sample; ROIs with Poisson error >10% were excluded. Incubations were amended with both ¹³C-bicarbonate and ¹⁵N-ammonium, and incorporation of both isotopes was evaluated. A T₀ sample consisting of PR10 sediment fixed immediately after addition of stable-isotope-labeled substrates was used as a common baseline for calculating atom percent excess (APE) and defining thresholds for detectable ¹³C and ¹⁵N enrichment, with immediate fixation minimizing biological incorporation of the added tracers. Daily measurements were adjusted by one-point correction to a check standard. Cells were classified as ¹³C- or ¹⁵N-active when their respective atom fraction exceeded the corresponding T₀ mean by >2 standard deviations, and APE was calculated as described by Pett-Ridge and Weber^68^.

### BONCAT-FACS, genome amplification, and sequencing

To recover DNA from anabolically active cell populations, sediment slurries from three PR07 horizons (0–10, 417–427 and 825–835 cmbsf) were analyzed by BONCAT-FACS-seq. Slurries were incubated with HPG for 10 weeks and subsequently preserved in glycine-TE buffer. Samples were processed using the cell extraction, on-filter DNA staining and BONCAT click-chemistry labeling workflow described above, with Alexa Fluor 405 azide dye (Vector Laboratories, Newark, CA, USA) used as the BONCAT fluorophore and with modifications to preserve cell integrity for sorting. Specifically, the sonication step used during standard cell extraction was omitted, and after labeling and washing, cells were gently resuspended from filters into 1× PBS containing 0.02% Tween 20 for flow cytometric sorting^15^.

Cell populations were sorted on a BD FACSDiscover S8 Cell Sorter (BD Biosciences, Franklin Lakes, NJ, USA) at the Scripps Research Flow Cytometry Core Facility. Instrument performance and optical alignment were verified using the facility’s standard quality-control and calibration procedures before sample acquisition and sorting. SYBR Green fluorescence was excited by the 488-nm blue laser and detected in the B3 channel (532/16-nm bandpass), while red fluorescence was excited by the 640-nm red laser and detected in the R2 channel (675/20-nm bandpass). These fluorescence parameters, together with forward and side scatter, were used to distinguish cellular particles from sediment particles and background events. Within the gated cell population, Alexa Fluor 405 fluorescence (V3 [460]-A) was used to separate HPG-labeled BONCAT-positive cells from the B-T fraction. An HPG-amended copper-omission click-chemistry control prepared from PR02 surficial sediment (0–10 cmbsf; 8.5 MPa) was used to characterize background Alexa Fluor 405 fluorescence and define the BONCAT-positive sorting threshold. A reagent-only PBS control containing the processing reagents in PBS but no sediment-derived cells was carried through BONCAT-FACS separately, and events within the sort gate were collected for downstream MDA and sequencing to identify reagent- and workflow-associated contaminants. Cells within the parent-cell gate were sorted into a BONCAT-positive fraction and a separate below-threshold fraction comprising cells that did not exceed the Alexa Fluor 405 fluorescence threshold used for BONCAT-positive sorting. The same gates were applied consistently across samples. Sorting was performed in purity mode into low-bind tubes containing PBS. Sample-specific acquired-event counts, sorted-cell counts, sequencing results and read-QC summaries are provided in Supplementary Table 10.

Because sorted-cell biomass was low, cells were lysed and genomic DNA was amplified using the REPLI-g Single Cell Kit (QIAGEN, Hilden, Germany; cat. no. 150343) according to the manufacturer’s single-cell protocol. Following cell lysis and DNA denaturation/neutralization, Phi29 polymerase-based multiple displacement amplification (MDA) was performed at 30 °C for 8 h, followed by polymerase inactivation at 65 °C for 3 min. The reagent-only PBS control was carried through MDA and sequencing alongside the sorted experimental fractions. Accordingly, BONCAT-FACS-seq profiles were interpreted as the taxonomic composition of recovered, detectably labeled cells rather than as quantitative estimates of total-community composition. Amplified DNA was submitted to the DNA Technologies and Expression Analysis Core at the UC Davis Genome Center for Illumina library preparation and sequencing on an Illumina NovaSeq X platform using 2 × 150 bp paired-end sequencing. Sequencing and read-QC summaries are provided in Supplementary Table 10.

### BONCAT-FACS-seq bioinformatic analysis

Raw BONCAT-FACS-seq reads were assessed using FastQC v0.12.1^69^ and adapter- and quality-trimmed using Trimmomatic v0.39^70^ with the parameters ILLUMINACLIP:TruSeq3-PE-2.fa:2:30:10, LEADING:3, TRAILING:3, SLIDINGWINDOW:4:15 and MINLEN:36. Post-trimming read quality was assessed using FastQC and summarized using MultiQC v1.19^71^. Taxonomic classification was performed using Kraken2 v2.14^72^ against a custom GTDB release 226 database downsampled to a maximum hash-table size of 128 GB using the Kraken2 --max-db-size option, with a classification confidence threshold of 0.05. Taxonomic abundances were estimated using Bracken v3.1^73^ with a read length of 150 bp and a minimum abundance threshold of 10 reads; genus-level profiles were used for downstream community-composition analyses. Genera detected in the sequenced reagent-only PBS control were removed. Relative abundances were recalculated from the retained Bracken-estimated reads after contaminant filtering. Bray–Curtis dissimilarity between each sorted fraction and its corresponding T₀ metagenomic community was calculated from the resulting genus-level relative-abundance profiles after normalization to unit sum.

For genome-resolved recruitment, quality-controlled paired-end reads were mapped against the Puerto Rico Trench MAG collection^38^ (NCBI BioProject PRJNA1334354) using Bowtie2 v2.5.4^74^ with the --very-sensitive-local setting, retaining a single best alignment per read (-k 1) and allowing insert sizes up to 2,000 bp. Primary alignments were processed using SAMtools v1.20^75^, with unmapped, secondary, and supplementary alignments excluded and a minimum mapping quality of 20. MAG recruitment and genome coverage were quantified using CoverM v0.7.0, retaining alignments with ≥95% nucleotide identity across ≥75% of the read. MAG–sample pairs with ≥20% genome breadth of coverage were classified as high-confidence recruitment; remaining non-zero recruitment was considered low-level.

### Phylogenomic analysis

JS1-affiliated MAGs and single amplified genomes (SAGs) from marine sediments were compiled from published studies, associated NCBI genome repositories, and genomes provided directly by principal investigators when available (Supplementary Table 14). Genome quality was assessed with CheckM v1^76^, and taxonomy was assigned with GTDB-Tk using GTDB release 226^77^ to confirm placement within the JS1 class-level lineage. Genomes were retained for phylogenomic analysis if they had ≥50% completeness and <10% contamination based on CheckM estimates. Genomes yielding all-gap marker alignments were excluded from downstream tree construction.

Environmental source metadata for each genome were obtained from the corresponding publication and, where available, associated NCBI metadata. Prior to comparative genomic analysis, sampling environments were manually classified into broader carbon/substrate regimes based on the reported depositional setting and substrate characteristics; genome-specific assignments and source metadata are provided in Supplementary Table 14.

Phylogenomic reconstruction was performed with the non-dereplicated MAG/SAG collection so that closely related genomes were retained and fine-scale lineage structure could be visualized. Conserved bacterial single-copy marker genes were identified and concatenated with GToTree v1.6.31 using the built-in bacterial HMM set in best-hit mode^78^. Maximum-likelihood phylogenies were inferred from the concatenated protein alignment using IQ-TREE^79^, with ModelFinder used to select the best-fit substitution model^80^ and branch support assessed with 1,000 ultrafast bootstrap replicates^81^. The final tree was visualized and annotated in iTOL v7.6^82^. Pairwise AAI was calculated from predicted protein sets using CompareM v0.1.2 to evaluate genome-relatedness within and between JS1 clades. AAI values of 65–95% and ≥95% were interpreted as approximate genus-like and species-like similarity ranges, respectively, following commonly used genome-relatedness thresholds^83^.

### Genome architecture, functional annotation, and physicochemical properties

Genome architecture metrics were calculated for the retained JS1 MAG/SAG set using CheckM v1.2.5 and Prodigal v2.6.3. Outputs included observed genome size, corrected genome size, GC content, predicted ORF count, and predicted ORF density^76,84^. Corrected genome size was calculated as observed genome size × 100 / (completeness − contamination), and ORF density was calculated as predicted ORFs per corrected Mbp. Predicted proteins were assigned to KOs using KOfamScan v1.3.0^85^.

Proteome physicochemical properties were calculated from Prodigal-predicted proteins using the Peptides package^86^. Genome-level proteome summaries included protein isoelectric point, acidic:basic residue ratio, charged residue fraction, hydrophobic residue fraction, and amino-acid composition. Genome architecture, KO-assignment, and proteome physicochemical metrics were summarized for non-hadal JS1 genomes and hadal-associated JS1 clades resolved in the phylogenomic analysis.

### Statistical analyses

Statistical analyses were performed in R v4.5.3, with multivariate analyses conducted using vegan v2.7-5^87^. BONCAT-positive proportions were measured in two incubation replicates per sediment horizon and pressure treatment and were therefore compared descriptively using replicate values, fold differences between treatment means, and PAI. The relationship between water depth and the mean *in situ*-pressure BONCAT-positive proportion in surficial sediments was tested using a two-sided Spearman rank correlation across the seven sites.

Cellular isotope-incorporation analyses were restricted to isotopically enriched (“active”) ROIs interpreted as cells because sediment background prevented reliable identification and enumeration of all unenriched cells in the ¹²C¹⁴N⁻ and ¹²C¹²C⁻ images. Cellular ¹⁵N atom percent excess distributions were compared between pressure treatments within each site using two-sided Wilcoxon rank-sum tests with Bonferroni correction for two comparisons. Because each site and pressure treatment comprised one nanoSIMS incubation, these tests compare cell-level distributions within incubations and do not represent incubation-level replication. Gini coefficients were calculated separately from ¹³C and ¹⁵N APE values of active cells using DescTools v0.99.60 to quantify cell-to-cell heterogeneity in isotope assimilation^31^.

Associations among geochemical variables, biomass-linked proxies, rarefied ASV richness and JS1 relative abundance were tested using two-sided Spearman rank correlations. Variables measured at different sediment depths were paired within sites using maximum-cardinality, minimum-total-offset one-to-one matching without replacement; matching tolerances and median depth offsets are reported in Supplementary Table 4. O₂ was linearly interpolated to target horizons and set to zero at and below the first measured zero. The 15 transect-wide correlations were adjusted together using the Benjamini–Hochberg procedure. Correlations between JS1 relative abundance and geochemical variables constituted a separate analysis family and were adjusted separately using the Benjamini–Hochberg procedure (Supplementary Table 13).

Group-wise comparisons among non-hadal JS1 genomes and the two hadal-associated clades were performed using pairwise Wilcoxon rank-sum tests, with *P* values adjusted for multiple comparisons using the Benjamini–Hochberg procedure (Supplementary Table 16). These tests were applied to genome architecture metrics, KO-assignment metrics and predicted proteome physicochemical properties. Because genome type and quality differed among JS1 genomic groups (Extended Data Fig. 5), covariate-adjusted linear models were used as robustness checks for selected genome architecture, KO-assignment and proteome physicochemical metrics. These models included JS1 group as the predictor of interest and CheckM-estimated completeness, CheckM-estimated contamination and genome type as covariates. Statistical outputs are reported in Supplementary Table 17.

Multivariate differences in KO-based functional profiles were assessed using Bray–Curtis dissimilarities calculated from KO copy densities normalized per 1,000 predicted proteins. Ordination was performed by principal coordinate analysis (PCoA), and group differences were tested using PERMANOVA with adonis2 in vegan, with additional models including carbon/substrate regime as a covariate. Differences in multivariate dispersion were evaluated using betadisper in vegan. Statistical outputs are reported in Supplementary Table 15.

## Acknowledgements

We thank the captain, crew, and scientific party of the R/V *Neil Armstrong* cruise AR64-02 for sample collection and shipboard support. We also thank undergraduate volunteers in the Bartlett laboratory at the Scripps Institution of Oceanography for assistance with sample processing and laboratory work. Cell extraction and enumeration were performed at the Kochi Institute for Core Sample Research, Japan Agency for Marine-Earth Science and Technology (JAMSTEC). Microscopy imaging was performed at the UC San Diego School of Medicine Microscopy Core, supported by NIH grants NS047101, OD030505, and OD036455. Part of this work was performed at nano@stanford RRID:SCR_026695. We thank Matthew Mills and Christie Jilly for assistance with nanoSIMS operation. Sequencing was carried out at the DNA Technologies and Expression Analysis Cores at the UC Davis Genome Center (RRID:SCR_017740), supported by NIH Shared Instrumentation Grant 1S10OD010786-01. Fluorescence-activated cell sorting was performed at the Scripps Research Flow Cytometry Core. We thank Gregory Henkes and Yang Gao for assistance with TOC and δ¹³C_org_ analyses.

## Funding

This work, including ship time, sample collection, analytical materials, and partial stipend support for M.D., was supported by the National Science Foundation (OCE-2049515). M.D. was also supported by a NASA Future Investigators in NASA Earth and Space Science and Technology (FINESST) award (80NSSC24K1832). L.W. acknowledges funding by a Minghua Zhang Faculty Career Catalyst Award. The funders had no role in study design, data collection and analysis, decision to publish, or preparation of the manuscript.

## Author contributions

M.D. designed the experiments, collected samples, performed laboratory experiments, analyzed and interpreted the data, prepared the figures, and wrote the manuscript. C.B. performed metagenomic analyses and contributed to data interpretation. A.H.F. performed nanoSIMS analyses and interpreted the data, with C.M.J.B. supervising nanoSIMS measurements. W.D.O. generated the total adenylate data. A.M.P. participated in shipboard sampling and laboratory work. Y.M. provided guidance on cell extraction methods. L.M.W. generated total organic carbon and δ¹³Corg data. D.G. performed dissolved geochemical measurements. R.A.P. generated the bathymetric map. A.J.S., I.A., and T.W. participated in shipboard porewater sampling, including sample selection and preservation for post-cruise geochemical analyses. B.S., S.R.S., and J.S.-C. participated in shipboard sampling and sample processing. Y.Z. participated in shipboard sediment sampling and sample curation that contributed to the molecular datasets used in this study. E.A.S. participated in shipboard porewater sampling, including sample selection and preservation for post-cruise geochemical analyses. D.C.S. participated in shipboard operations and sample collection. A.E.D. supervised the nanoSIMS analyses. J.S.B. provided computational resources. All authors reviewed and edited the manuscript and approved the final version. D.H.B. and S.D. conceived and supervised the study and secured funding.

## Competing interests

The authors declare no competing interests.

## Data availability

The sequencing data generated in this study have been deposited in the NCBI Sequence Read Archive under BioProject accession PRJNA1541392. The data will be made publicly available upon publication and are available to reviewers upon request.

## Code availability

Custom code used for bioinformatic and statistical analyses in this study is publicly available on GitHub at https://github.com/migdesmarais/Puerto-Rico-Trench-Piezophily.

## Extended Data Figure Legends

**Extended Data Fig. 1.**
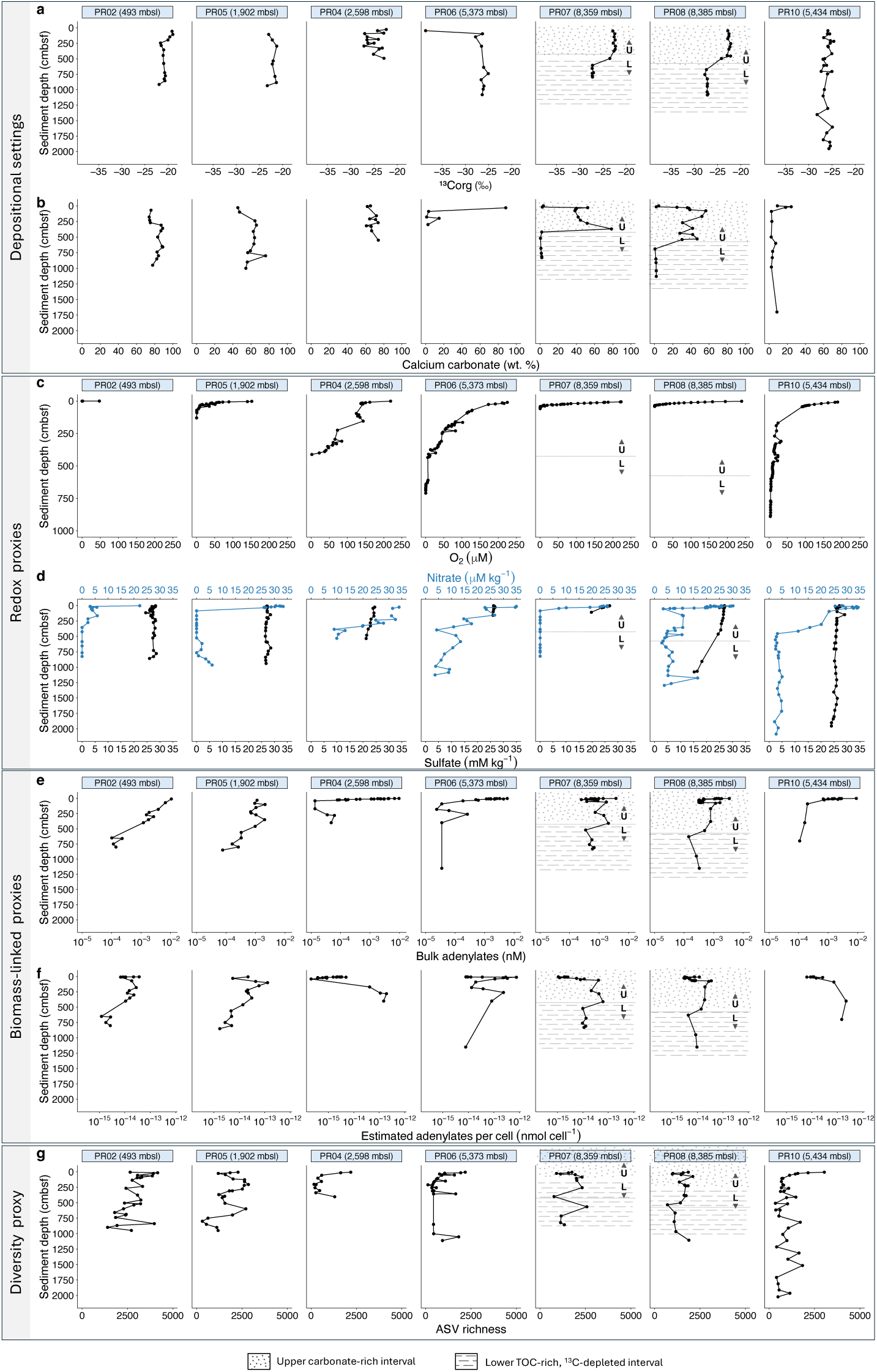
Geochemical, redox, biomass-linked and diversity profiles across the Puerto Rico Trench transect. a,b, δ¹³Corg (‰) and calcium carbonate (CaCO3; wt.%) profiles. c,d, Oxygen (µM), nitrate (µM kg^-1^) and sulfate (mM kg^-1^) profiles. e, Total adenylate concentration (ATP + ADP + AMP; nM). f, Estimated adenylate content per cell (nmol cell⁻¹). g, Rarefied amplicon sequence variant (ASV) richness. Dashed lines mark the boundary between the upper carbonate-rich (U) and lower TOC-rich, ¹³C-depleted (L) intervals at PR07 and PR08.

**Extended Data Fig. 2.**
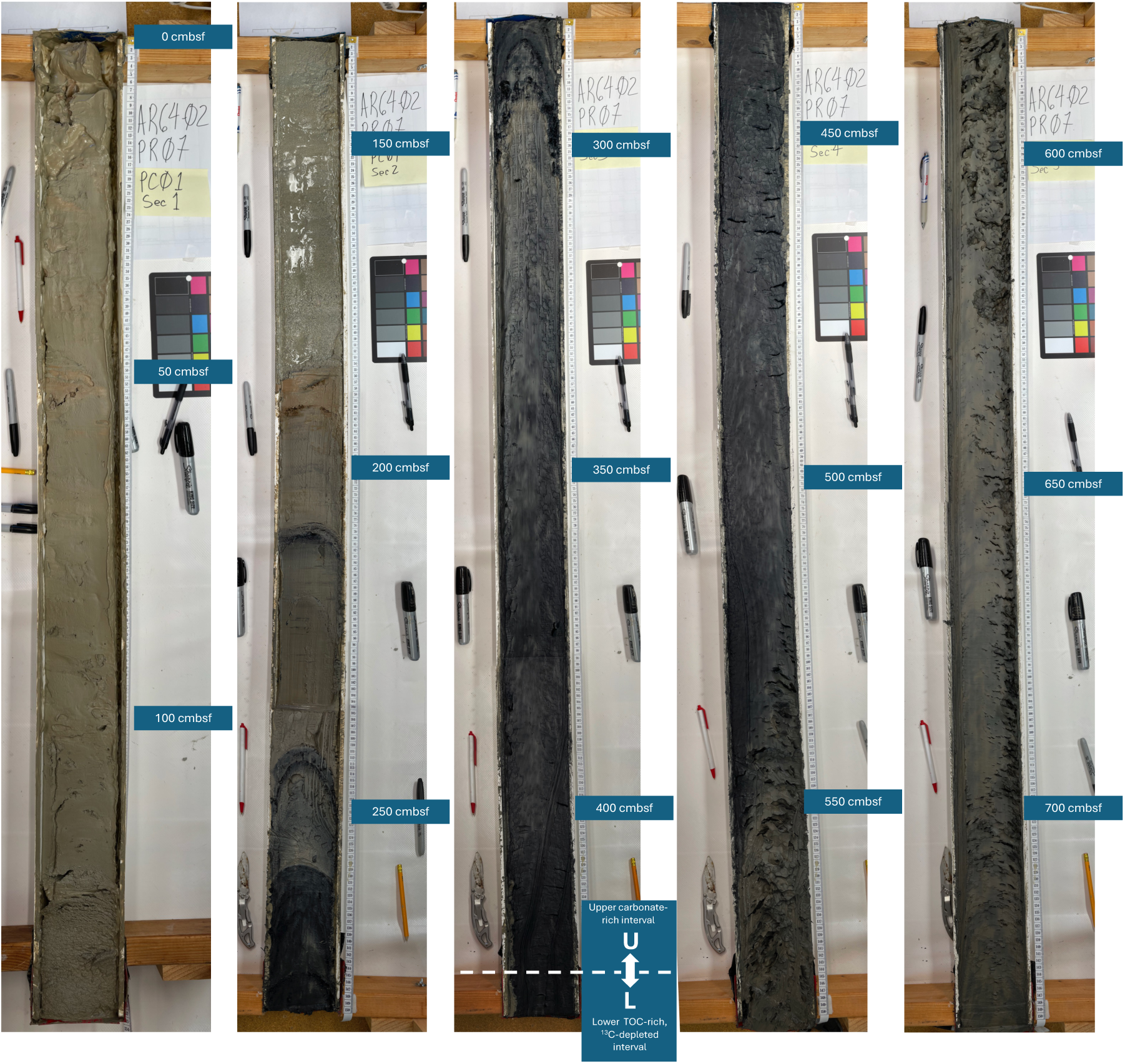
Lithological transition in PR07 trench-axis sediments. Split-core images showing the transition from the lighter-colored upper carbonate-rich interval (U) to the darker-colored lower TOC-rich, ¹³C-depleted interval (L). The boundary between intervals occurs at approximately 425 cmbsf.

**Extended Data Fig. 3.**
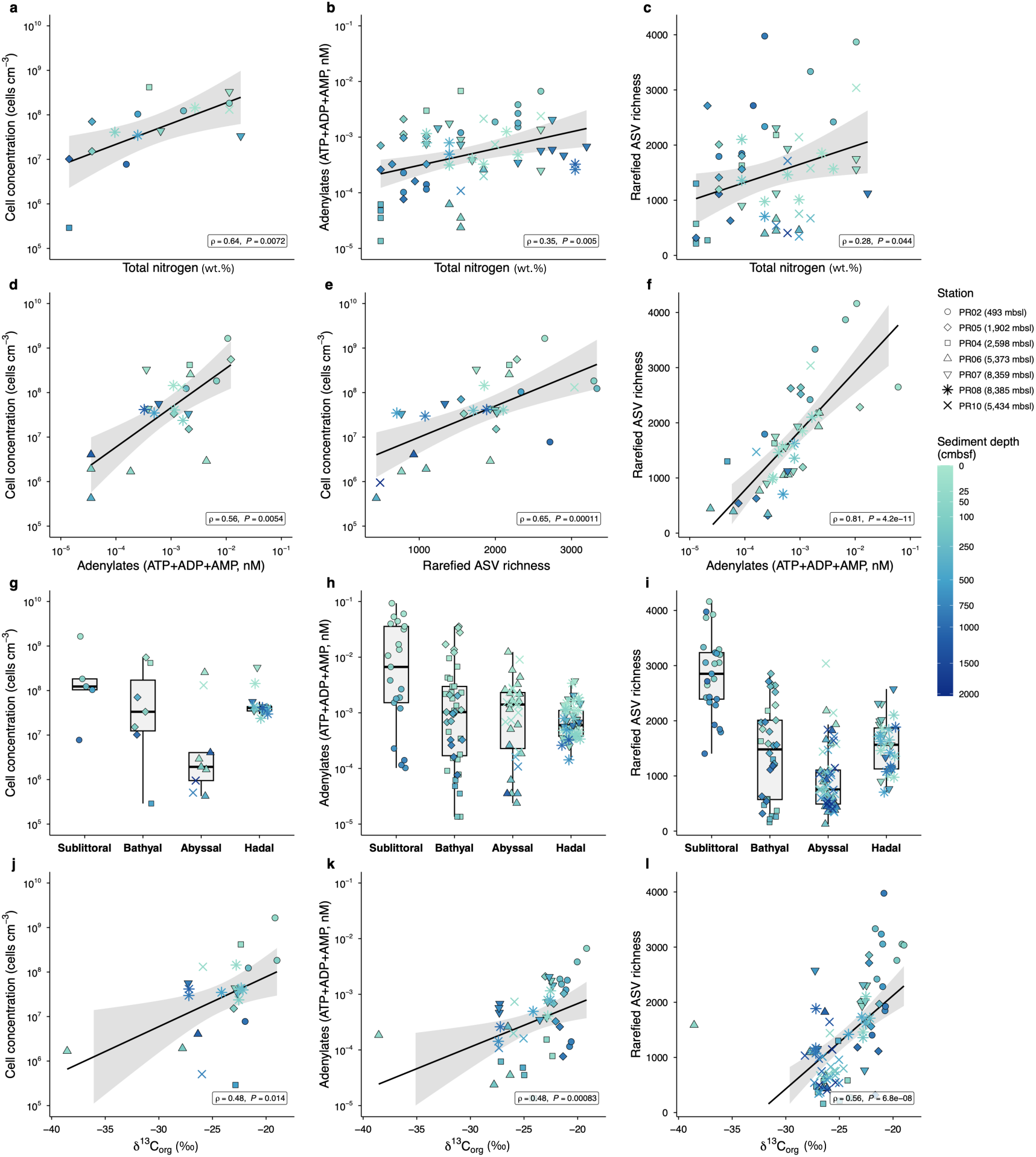
Relationships among organic matter properties and microbial proxies across the Puerto Rico Trench transect. **a–c**, Relationships of total nitrogen content (TN; wt.%) with cell concentration (cells cm^-3^), total adenylate (ATP + ADP + AMP; nM) and rarefied amplicon sequence variant (ASV) richness, respectively. **d–f,** Pairwise relationships among these microbial proxies. **g–i,** Distributions of each proxy across depositional settings. **j–l,** Relationships of δ¹³C_org_ (‰) with cell concentration, total adenylate and rarefied ASV richness, respectively. Black lines and shading show fitted relationships and 95% confidence intervals. Insets report Spearman ρ and *P* values.

**Extended Data Fig. 4.**
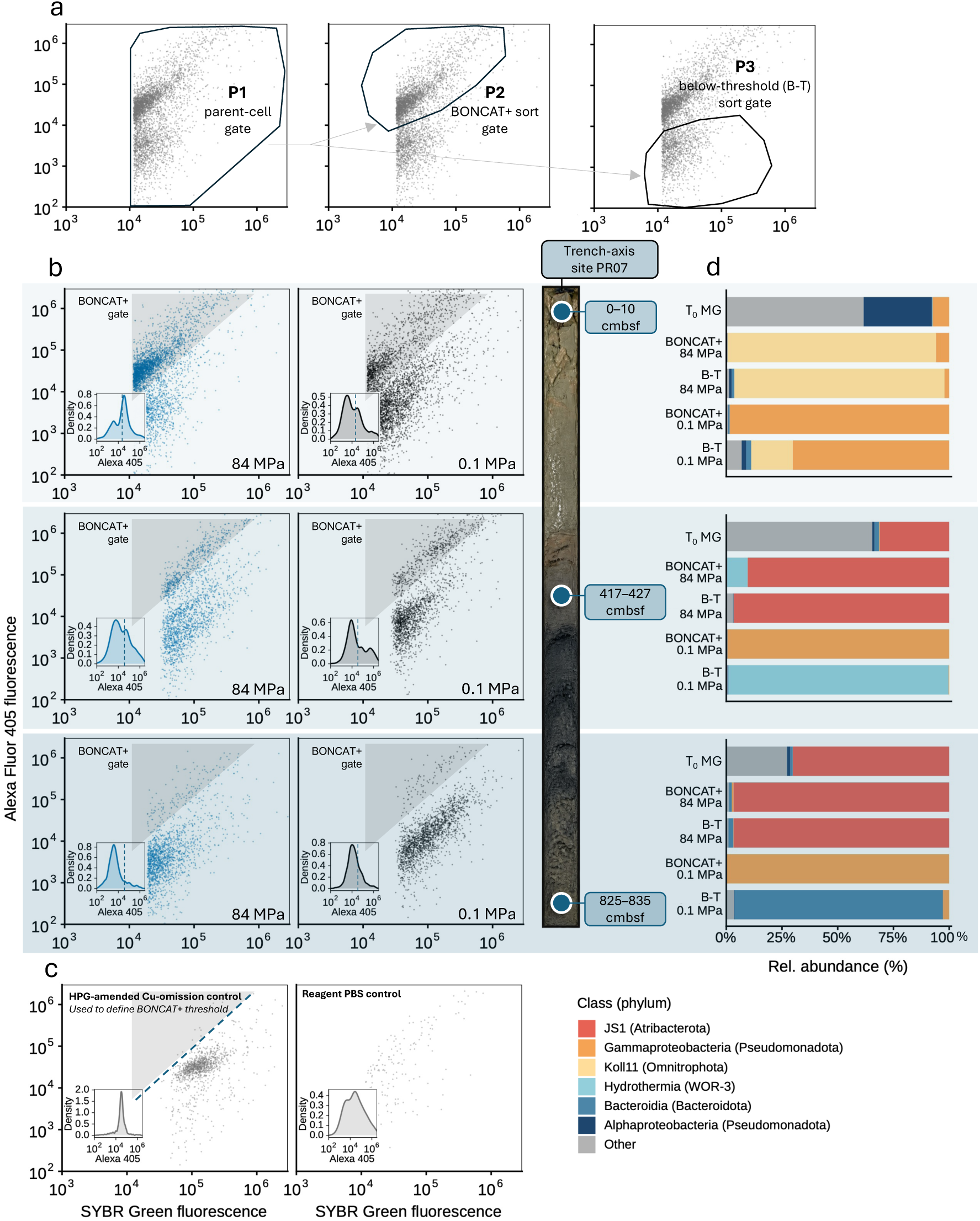
BONCAT-FACS gating and taxonomic composition of sorted PR07 sediment communities. **a**, Sequential FACS gating of parent cells (P1), BONCAT-positive cells (P2) and below-threshold (B-T) cells (P3). **b,** SYBR Green and Alexa Fluor 405 fluorescence during sorting of PR07 sediments at 0–10, 417–427 and 825–835 cmbsf incubated under atmospheric and *in situ* pressure; shading indicates the BONCAT-positive gate, and insets show Alexa Fluor 405 distributions. **c,** HPG-amended copper-omission control used to define the BONCAT-positive fluorescence threshold and reagent-only PBS control. **d,** Class-level taxonomic composition of T₀ metagenomes and BONCAT-positive and below-threshold sorted fractions across sediment depths and pressure treatments.

**Extended Data Fig. 5.**
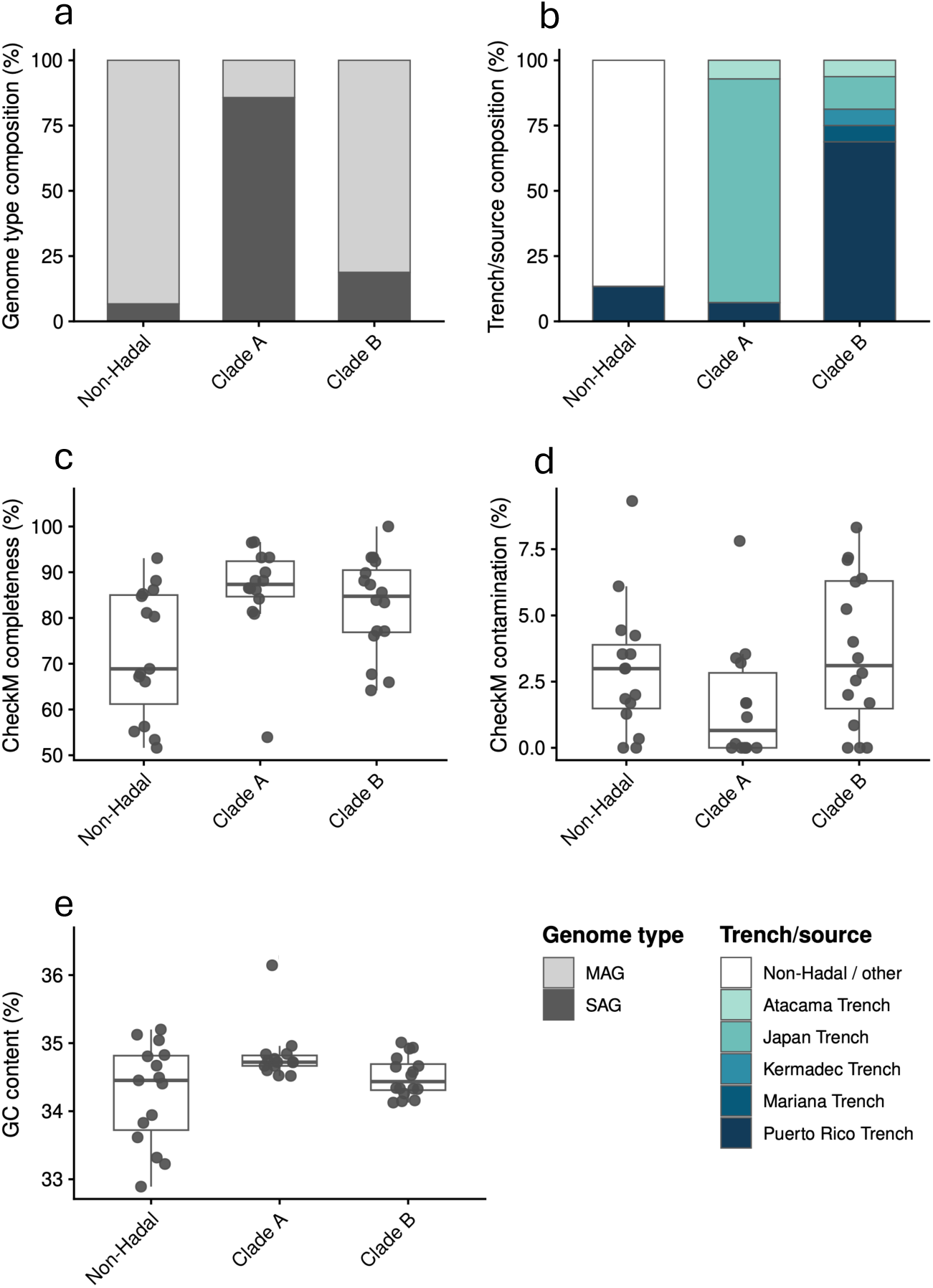
Composition and genome properties of the JS1 genome collection. **a**, MAG and SAG composition among non-hadal JS1 and hadal clades A and B. **b,** Trench and source composition of each genomic group. **c–e,** CheckM completeness (%), contamination (%) and GC content (%), respectively. Points represent individual genomes; boxes show medians and interquartile ranges.

**Extended Data Fig. 6.**
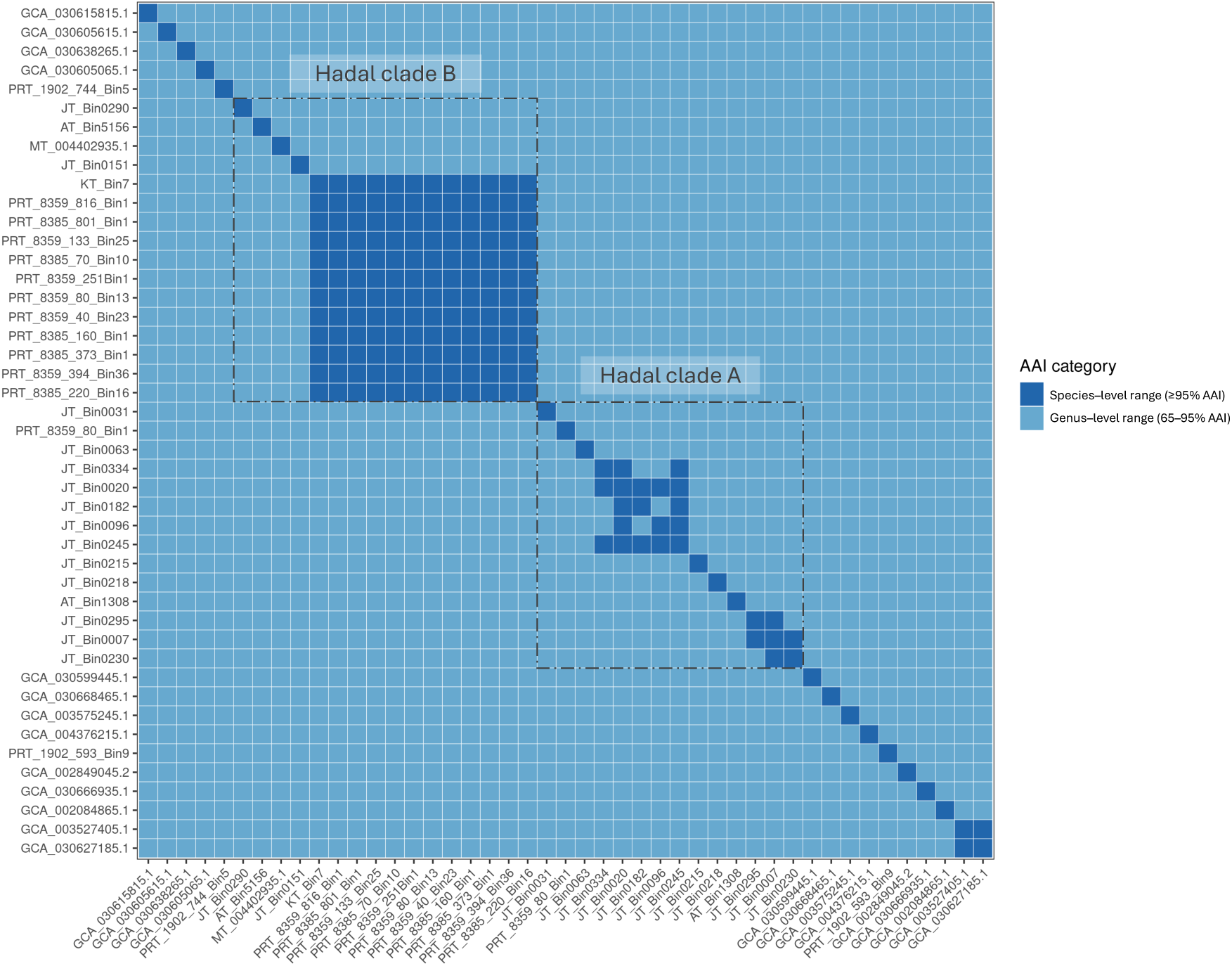
Pairwise average amino acid identity among JS1 genomes. Heatmap of pairwise average amino acid identity (AAI), with colors indicating genus-level (65–95% AAI) and species-level (≥95% AAI) ranges. Boxes delineate hadal clades A and B.

**Extended Data Fig. 7.**
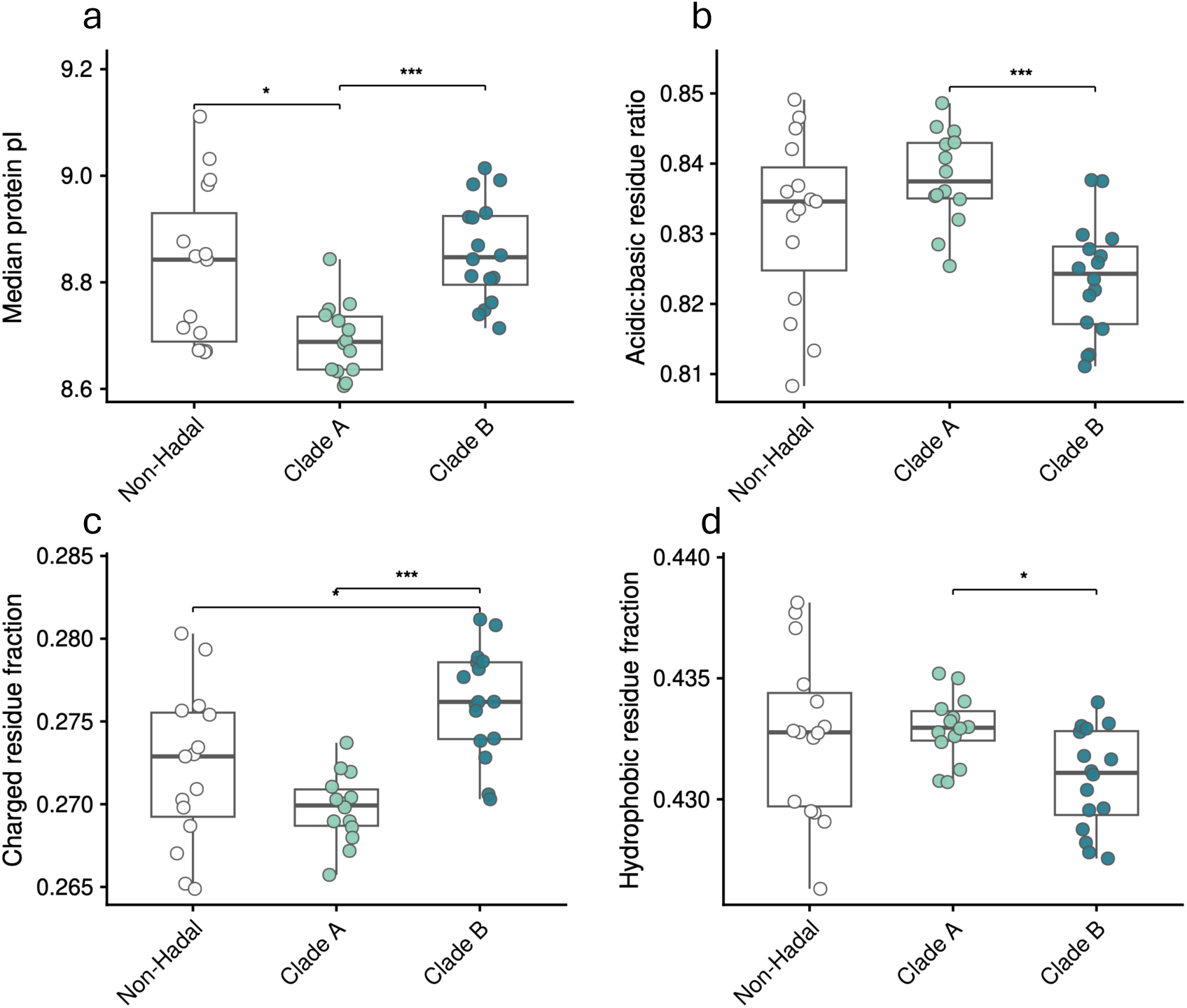
Proteome physicochemical properties across JS1 genomic groups. **a**, Median protein isoelectric point (pI). **b,** Acidic-to-basic residue ratio. **c,** Charged-residue fraction. **d,** Hydrophobic-residue fraction. Points represent individual genomes; boxes show medians and interquartile ranges. Pairwise Wilcoxon rank-sum tests with Benjamini–Hochberg-adjusted *P* values (\**P* < 0.05, \*\**P* < 0.01, \*\*\**P* < 0.001).

## Notes

### Competing Interest Statement

The authors have declared no competing interest.

### Summary of Updates

Updated manuscript to reflect the version submitted to Nature Microbiology, including minor text revisions, reorganization, and shortening of the main text.

